# 3’ Nucleotide Asymmetry Directs miRNA Strand Selection

**DOI:** 10.1101/2025.10.15.682714

**Authors:** Jeffrey C. Medley, Sumire Kurosu Moriya, Huiwu Ouyang, Heather Crawshaw, Sarah Y. Zhang, Ganesh Panzade, Will J. Sydzyik, Joel T. Sydzyik, Mira Bhandari, Christopher M. Hammell, Anna Zinovyeva

## Abstract

Accurate microRNA (miRNA) strand selection is essential for defining the regulatory landscape of the miRNA-induced silencing complex (miRISC). While 5′ nucleotide identity and duplex thermodynamics have been proposed to bias strand choice, these features cannot fully explain in vivo strand preferences. Here, we uncover a conserved and previously unrecognized role for 3′ nucleotide asymmetry in directing miRNA strand selection in *Caenorhabditis elegans* and human cells. A favorable 3′ terminal nucleotide on the passenger strand promotes selective loading of the opposing guide strand into miRISC, revealing a cooperative interplay between 5′ and 3′ terminal asymmetries that ensures precise strand discrimination. These findings establish a unified, evolutionarily conserved mechanism for miRNA duplex sorting and expand the fundamental rules governing small RNA biogenesis.

## Introduction

The biogenesis of miRNAs is integral to their biological activity, as the precise sequence of mature miRNAs must be preserved to maintain sequence-specific interactions with mRNA targets (*1, 2*). Primary miRNA (pri-miRNA) transcripts are processed in the nucleus by the Microprocessor complex to produce stem-loop miRNA precursors (pre-miRNA) (*3*–*6*). Following nuclear export, Dicer cleaves pre-miRNAs to generate partially double-stranded ∼22 nucleotide duplexes containing characteristic two-nucleotide 3’ overhangs (*7*–*12*). The miRNA duplex is subsequently loaded into an Argonaute protein, which selectively anchors a single, mature strand to initiate formation of the miRNA induced silencing complex (miRISC) (*13, 14*). Although the strands originating from either the 5’ (5p) or 3’ (3p) arms of the pre-miRNA are competent for Argonaute loading (*15*–*17*), asymmetric strand selection usually leads to a dominant guide strand and a less abundant passenger strand that is rapidly degraded after release from Argonaute (*18*). For some miRNAs, strand selection occurs in a more symmetrical fashion where both strands appear to function as mature miRNAs (*19, 20*). Differences in miRNA strand usage have also been observed in different cellular, tissular and developmental contexts, suggesting that strand selection is a regulated process (*19, 21*–*24*). As miRNA-dependent gene regulation is sequence dependent, alternative miRNA strand choice is expected to lead to broad changes in target selection. Consistently, aberrant miRNA strand selection has also been reported in several human diseases including cancers, although the mechanisms leading to alternative strand selection in human disease are not fully understood (*18*).

Argonaute plays a direct role in strand selection and favors loading the duplex end containing a favorable 5’ nucleotide (U>A>C>G) and lower thermodynamic stability in vitro (*16, 17, 25, 26*). The 5’ nucleotide of the guide strand is embedded within a pocket in the MID domain of Argonaute, which has a strong preference for uracil (*13,14, 25*). Several residues within the Argonaute MID and PIWI domains directly contact the phosphate backbone of the guide strand (*13, 14*) and may serve as a sensor for thermodynamic asymmetry (*17*). The twin-drive model postulates that miRNA strand selection can be predicted by modeling the relative effects of 5’ nucleotide identity and thermodynamic asymmetry (*17*). However, this model fails to explain strand selection of ∼25% of miRNAs that have unfavorable 5’ nucleotides and more stable 5’ ends (*18*), suggesting that additional asymmetries influence miRNA strand selection in vivo.

Here, we used genome editing to alter endogenous *C. elegans* miRNAs and evaluated the relative contributions of 5’ nucleotide identity and thermodynamic asymmetry in vivo. Consistent with previous findings, we show that a 5’ uracil is favorable for stand choice, and that unpaired 5’ nucleotides are strongly selected regardless of nucleotide identity. However, we found that duplexes with symmetrical 5’ nucleotides and thermodynamic end stabilities underwent asymmetric strand selection, suggesting that additional features outside of the 5’ end contribute to strand choice. We identified a strong bias for cytosine at the 3’ end of passenger strands in *C. elegans*. Mutating the 3’ nucleotide of symmetrical duplexes was sufficient to reverse strand selection, suggesting that 3’ nucleotide identity influences miRNA strand asymmetry in vivo. In duplexes with asymmetric 5’ ends, 3’ nucleotide asymmetry had a lesser effect on strand choice, indicating that various nucleotide asymmetries coordinately drive miRNA strand selection. By expressing exogenous miRNA variants in human HEK293T cells, we show that 3’ nucleotide asymmetry plays a conserved role in miRNA strand selection, although a 3’ cytosine on passenger strands was less preferred in human cells. Collectively, our results support a model where nucleotide asymmetries on both strands of miRNA duplexes promote asymmetric strand selection.

## Results

### Characterization of *C. elegans* miRNA duplexes

To better understand how 5’ nucleotide identity and thermodynamic asymmetries are linked to miRNA strand selection, we profiled the duplex characteristics of *C. elegans* miRNAs (Table S1). The preferred guide strand is expected to have a favorable 5’ nucleotide and lower 5’ end stability than the corresponding passenger strand (Fig. 1A). Consistent with previous observations (*27*), the majority of *C. elegans* guide strands have a 5’ uracil, whereas passenger strands lack an obvious 5’ nucleotide bias (Fig. 1B). To examine thermodynamic asymmetry, we used free energy values of *C. elegans* miRNA duplex ends from a previous study (*28*). As expected, the average stability of duplex ends corresponding to guide strands was lower than passenger strands (Fig. 1C, Table S1). We observed thermodynamic asymmetry when considering the first through fourth terminal nucleotides, with the largest asymmetry observed for three terminal nucleotides (Fig. 1C). Thus, strand selection of *C. elegans* miRNAs is correlated to 5’ nucleotide identity and thermodynamic asymmetry.

**Fig 1.**
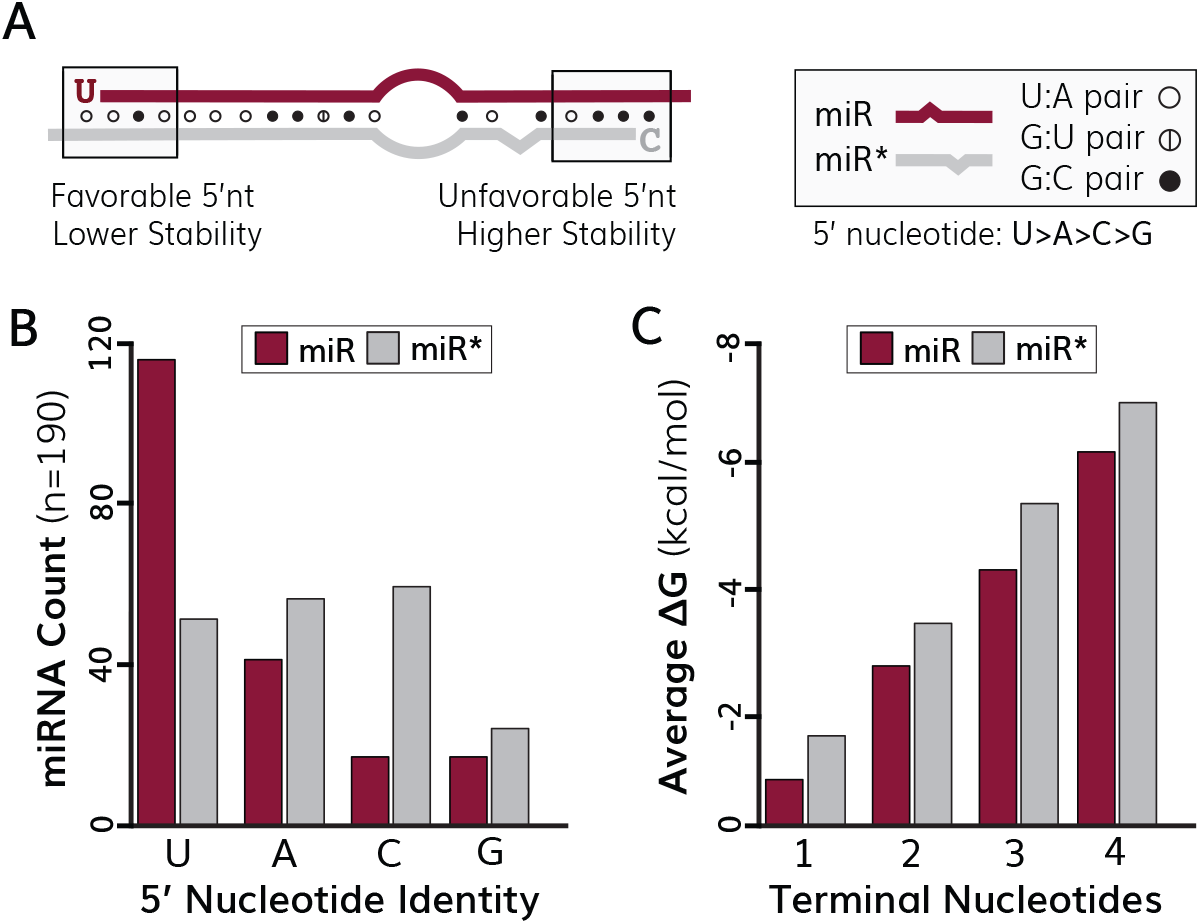
Duplex characteristics of C. elegans miRNAs. (A) Duplex features associated with miRNA strand selection. The guide (miR) end of typically contains a favorable 5’ nucleotide and lower thermodynamic stability than the passenger (miR^*^) duplex end. The guide strand is shown in maroon, and the passenger strand is shown in gray. U:A pairs are illustrated by an open circle, G:U wobble pairs are indicated by an open circle with a vertical line and G:C pairs are represented by closed circles. The sizes of bulges and mismatched nucleotides are drawn to scale. (B) Quantification of 5’ nucleotide identity for *C. elegans* miRNA guide (maroon) and passenger strands (gray). n is given as the number of miRNA duplexes quantified (C) Quantification of the free energy of miRNA duplex ends corresponding to guide (maroon) and passenger (gray) duplex ends. Free energy values were calculated by in silico unwinding *(28)* of the terminal 1:4 nucleotides of each duplex end.

### A favorable 5’ nucleotide is not sufficient to drive miRNA strand selection in vivo

To address whether 5’ nucleotide identity and thermodynamic asymmetry directly influence miRNA strand choice in vivo, we used genome editing to alter the duplex properties of endogenous *C. elegans* miRNAs. We first examined a series of mutations in the *mir-58a, mir-1* and *mir-2* miRNAs that primarily impacted the identity of 5’ nucleotides (Fig. 2, Table S2). Both strands of the miR-58 duplex have a favorable 5’ uracil, whereas only the guide strands of miR-1 and miR-2 have a 5’ uracil (Fig. 2). We used RNAcofold to determine how each miRNA variation affected the thermodynamic stability of the terminal three nucleotides for each duplex end (*28*) (Table S3). For all three miRNAs, the guide duplex end has a lower stability than the passenger duplex end (Fig. 2). Based on predictions from the twin-drive model (*17*), the four miR-58 mutations (M1-M4) should switch strand selection, the miR-1-M1 mutation should retain normal strand choice, and the miR-2-M1 mutation should choose both duplex strands indiscriminately leading to equal accumulation of both strands (Table S4).

**Fig 2.**
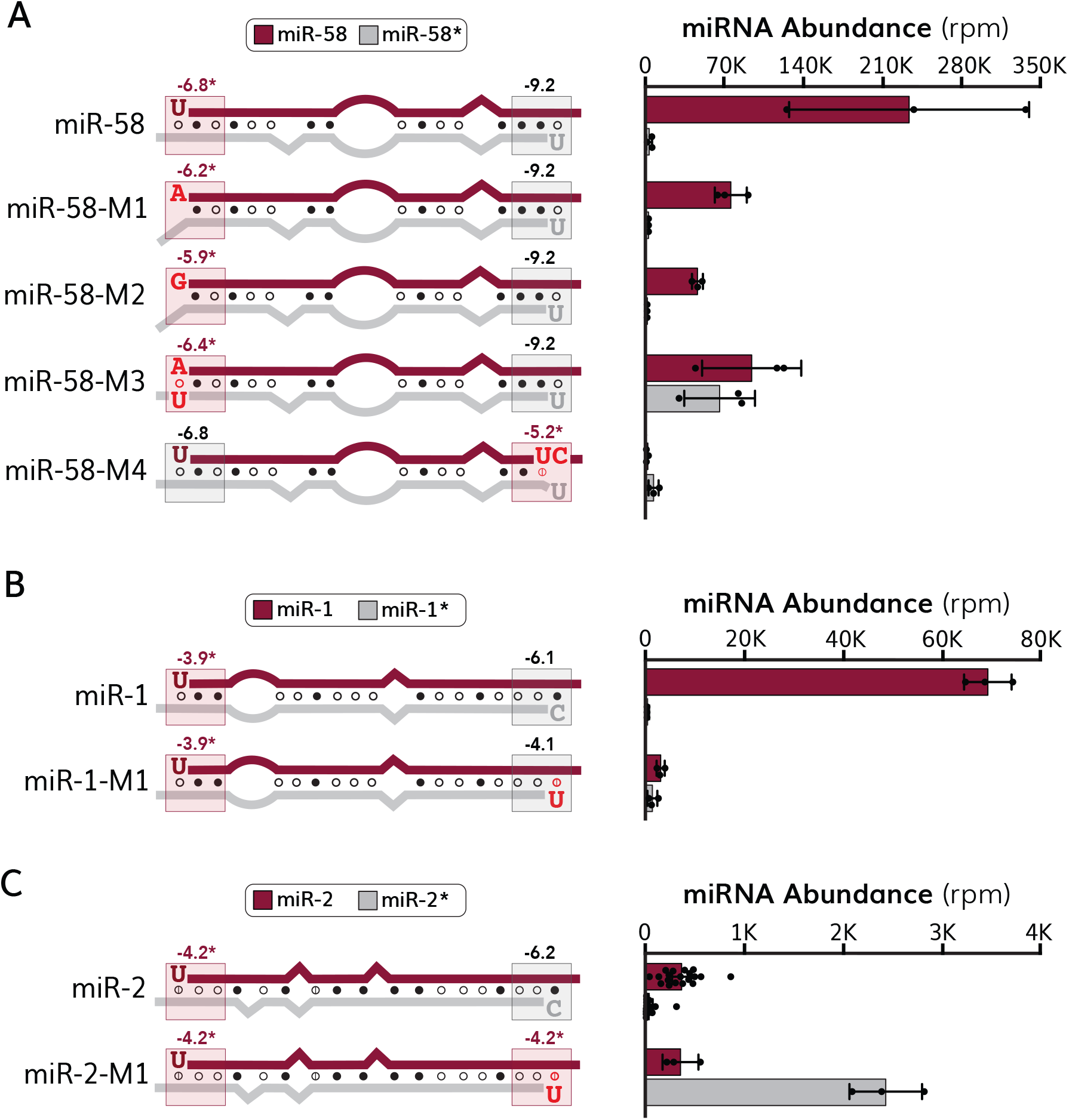
5’ nucleotide identity influences miRNA strand selection in vivo. (Left) Schematic representations of (A) miR-58, (B) miR-1 and (C) miR-2 duplex mutations that affect 5’ nucleotide identity. (A-C) The guide strand is shown in maroon, and the passenger strand is shown in gray. Mutated nucleotides are indicated by red text. U:A pairs are illustrated by an open circle, G:U wobble pairs are indicated by an open circle with a vertical line and G:C pairs are represented by closed circles. Free energy values (kcal/mol) of the terminal three nucleotides (boxed) are shown on each duplex end. The less stable duplex end is marked with an asterisk and highlighted in a light red box. RNAcofold was used to predict duplex structures at 20°C. (Right) Quantification of small RNA sequencing results for (A) miR-58, (B) miR-1 and (C) miR-2. Data are presented as mean ± standard deviation, with each dot representing quantification from a single library using the NEXTFLEX v3 Small RNA Sequencing Kit. Maroon bars correspond to miRNA guide strands and gray bars represent miRNA passenger strands. Normalization was performed by dividing raw read counts by total mapped miRNA reads and multiplying by one million to acquire reads per million (rpm) values.

To investigate how miRNA mutations affect strand selection, we performed small RNA sequencing in L4-staged animals (Tables S5 and S6), a developmental stage where miR-58, miR-1 and miR-2 have relatively high expression (*29*). The miR-58-M1 and miR-58-M2 mutations change the 5’ uracil of the guide strand to less favorable nucleotides (adenine and guanine respectively), which results in the 5’ nucleotide of the guide strand becoming unpaired, thereby modestly reducing duplex end stability (Fig. 2A). In the miR-58-M3 mutation, we introduced a compensatory mutation in the passenger strand that restores base pairing to the 5’ adenine variant (Fig. 2A). Our analysis revealed that miR-58-M1 and miR-58-M2 retain normal strand choice, while miR-58-M3 accumulated both duplex strands at near equal levels (Fig. 2A). As the primary difference between the miR-58-M1 and miR-58-M3 mutations is the base-pairing status of the 5’ nucleotide, these results suggest that unpaired 5’ nucleotides are highly favorable for strand selection and can override a preferrable 5’ nucleotide on the other strand. Consistently, the miR-58-M4 mutation, which alters the 3’ end of the guide strand to disrupt base-pairing of the 5’ nucleotide on the passenger strand, appeared to select the passenger strand although total miR-58 levels were reduced (Fig. 2A).

Both miR-1-M1 and miR-2-M1 mutations change a terminal C:G pair on the passenger end of the duplex into a U:G wobble pair, such that both duplex strands have a symmetrical 5’ uracil and similar thermodynamic end stabilities (Figs. 2B and 2C). For the miR-1-M1 mutation, we observed some accumulation of passenger strand, however, the miR-1-M1 mutation also appeared to affect mature miR-1 levels (Fig. 2B). On the other hand, we found that the miR-2-M1 mutation caused a reversal of miR-2 strand selection (Fig. 2C). Given that the miR-2-M1 mutation has symmetrical 5’ nucleotide identity and thermodynamic end stability (Fig. 2C), this suggests that additional factors likely drive asymmetric strand selection of miR-2. Notably, our data show that the twin-drive model poorly predicted the actual strand preference of these miRNA mutants (Table S4). We conclude that 5’ nucleotide identity directly influences miRNA strand selection in vivo, although favorable 5’ nucleotides are not sufficient to drive asymmetric strand choice, particularly when the 5’ nucleotide of one strand is unpaired.

### Thermodynamic asymmetry is largely dispensable for miRNA strand choice

We next examined mutations affecting thermodynamic asymmetry of *mir-58a, mir-1* and *mir-2* (Fig. 3). The miR-58-M5, miR-58-M6 and miR-58-M7 mutations reduce thermodynamic end stability without altering 5’ nucleotide identity (Fig. 3A). The twin-drive model (*17*) predicted each of these mutations, except for miR-58-M6 and miR-1-M2, to switch strands (Table S4). Small RNA sequencing of L4-staged animals (Tables S5 and S6) showed that the symmetrical miR-58-M5 and miR-58-M6 mutants asymmetrically selected the canonical miR-58 guide strand (Fig. 3A). Furthermore, while miR-58-M7 mutants accumulated some passenger strands, they did not switch strand preference, despite the passenger duplex end having lower stability (Fig. 3A). This observation suggested factors outside 5’ nucleotide identity and thermodynamic asymmetry promote strand selection of miR-58.

**Fig 3.**
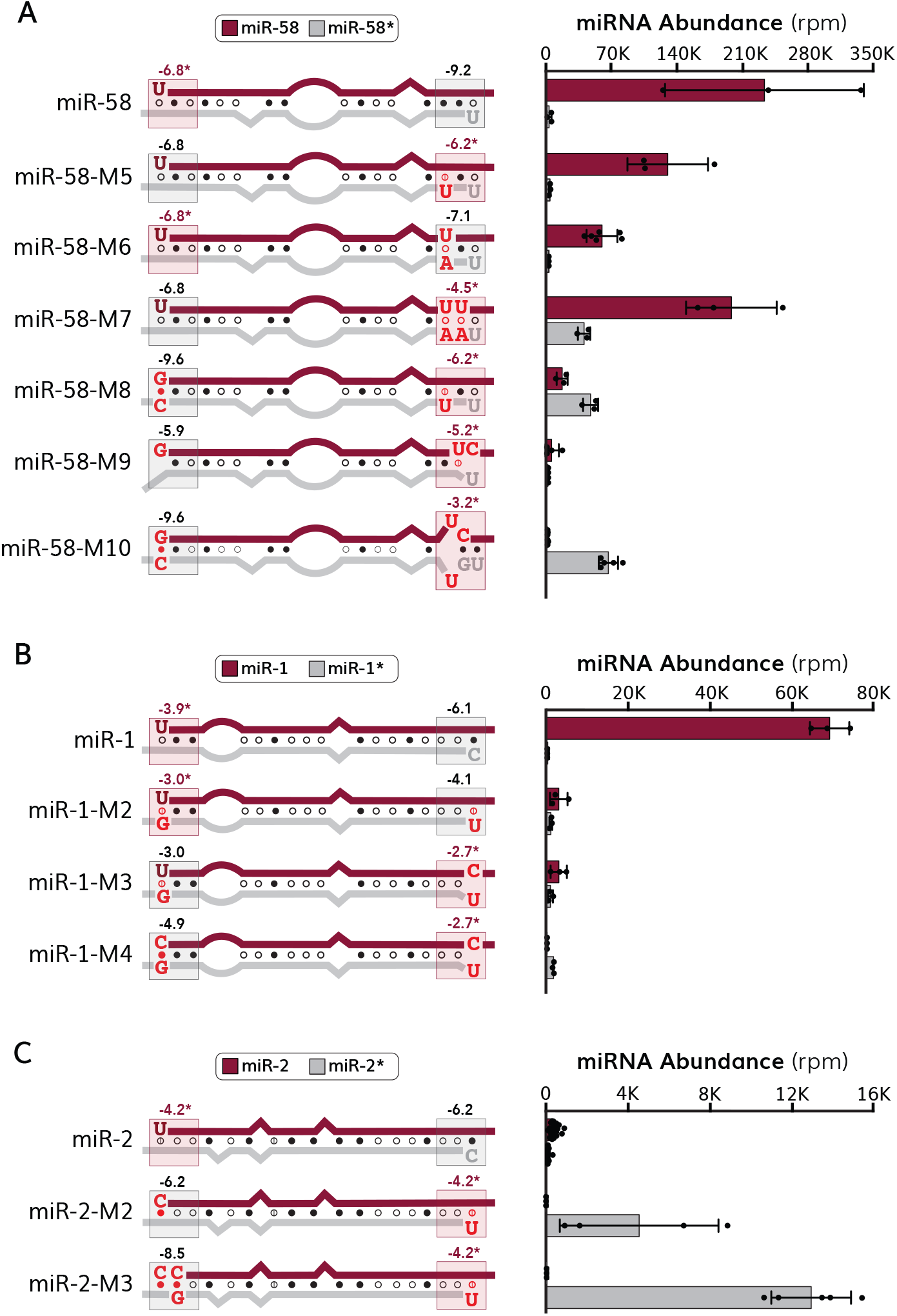
Thermodynamic asymmetry is dispensable for miRNA strand selection. (Left) Schematic representations of (A) miR-58, (B) miR-1 and (C) miR-2 duplex mutations that affect the relative thermodynamic stability of each duplex end. (A-C) The guide strand is shown in maroon, and the passenger strand is shown in gray. Mutated nucleotides are indicated by red text. U:A pairs are illustrated by an open circle, G:U wobble pairs are indicated by an open circle with a vertical line and G:C pairs are represented by closed circles. Free energy values (kcal/mol) of the terminal three nucleotides (boxed) are shown on each duplex end. The less stable duplex end is marked with an asterisk and highlighted in a light red box. RNAcofold was used to predict duplex structures at 20°C. (Right) Quantification of small RNA sequencing results for (A) miR-58, (B) miR-1 and (C) miR-2. Data are presented as mean ± standard deviation, with each dot representing quantification from a single library using the NEXTFLEX v3 Small RNA Sequencing Kit. Maroon bars correspond to miRNA guide strands and gray bars represent miRNA passenger strands. Normalization was performed by dividing raw read counts by total mapped miRNA reads and multiplying by one million to acquire reads per million (rpm) values.

The more egregious miR-58-M8, miR-58-M9 and miR-58-M10 mutations reduce the stability of the passenger duplex end while also introducing an unfavorable 5’ nucleotide to the guide strand (Fig. 3A). The miR-58-M8 and miR-58-M10 mutations both reversed strand selection, demonstrating that there is not a barrier to selecting the miR-58 passenger strand (Fig. 3A). While the miR-58-M9 mutation did not appear to switch strands, we observed a substantial reduction in mature miR-58 levels (Fig. 3A), which was associated with a significant increase in *pri-mir-58* levels, suggesting ineffective processing of *pri-mir-58* (Fig. S1A). We did not observe significant changes in the levels or strand selection of the *mir-58* family members *mir-80, mir-81* or *mir-82* (*30*), supporting our observations are specific to *mir-58* (Fig. S2).

The miR-1-M2, miR-1-M3 and miR-1-M4 mutations affect thermodynamic asymmetry while also introducing a 5’ uracil to the passenger strand; the 5’ nucleotide of the miR-1 guide strand is also mutated to a less favorable cytosine in miR-1-M4 mutants (Fig. 3B). We found that the miR-1-M2 and miR-1-M3 mutations both preferred selection of the canonical miR-1 guide strand, while miR-1-M4 asymmetrically selected the passenger strand (Fig. 3B). We observed a reduction in mature miR-1 levels in each of the miR-1 mutants that we generated in this study (Figs. 2B and 3B), possibly due to decreased *mir-1* expression as *pri-mir-1* levels were significantly reduced in miR-1-M4 mutants (Fig. S1B). The miR-2-M2 and miR-2-M3 mutations incorporate a favorable 5’ uracil to the miR-2 passenger strand, while also introducing cytosines to the 5’ end of the miR-2 guide strand that increase the stability of the guide duplex end (Fig. 3C). The miR-2-M2 and miR-2-M3 mutations both resulted in highly asymmetric selection of the miR-2 passenger strand and appeared to increase mature miR-2 levels (Fig. 3C). We observed modest effects on *pri-mir-2* levels in miR-2 mutants, suggesting that any effects on mature miR-2 levels likely occur downstream of miRNA biogenesis (Fig. S1C). Notably, the twin-drive model poorly predicted the strand preference of most miRNA mutants (Table S4, Figure S3), suggesting additional factors contribute to strand selection in vivo. Absolute quantification of select miRNA mutants agreed with our small RNA sequencing analysis (Fig. S4). Further, we observed phenotypes consistent with loss of guide strand activity in strand switching mutants (Fig. S5). In the *mir-58* family mutant background, the miR-58-M8, miR-58-M9 and miR-58-M10 mutations caused a reduction in adult body size, consistent with loss of *mir-58* function (*30*) (Fig. S5A-B). Similarly, our *mir-1* and *mir-2* mutants showed resistance to levamisole and aldicarb respectively, as previously reported for loss of *mir-1* (*31*) and *mir-2* (*32*) function (Fig. S5C-D).

To address whether miRNA biogenesis was affected in miRNA mutants, we quantified how each mutation affected isomiR populations. With a few exceptions, most of the miR-58 variants did not significantly affect the distribution of isomiRs for guide or passenger strands compared to controls, suggesting that miRNA mutations do not affect production of the expected mature miRNA (Fig. S6, Tables S7 and S8). In miR-58-M4, miR-58-M9 and miR-58-M10 mutants, a four-base truncation of the 3’ end of the guide strand was more abundant than the canonical guide sequence without affecting isomiRs for passenger strands (Fig. S6, Tables S7 and S8). Each of these three mutations contain the same CA:UC mutation towards the 3’ end of the guide strand (Table S2), which may impact processing of these mutants, as we observed for miR-58-M9 (Fig. S1A), or could adversely affect the stability of the 3’ end of the miRNA within the Argonaute PAZ domain (*13, 14, 33, 34*) thereby leading to trimming of exposed 3’ ends (*35, 36*). Similarly, the miR-1-M4 mutation accumulated a one-base truncation of the 3’ end of the canonical guide strand without affecting passenger strand isomiRs, while isomiR populations appeared unaffected for other miR-1 mutants (Fig. S8A-B, Table S9). The isomiR populations of miR-2 appeared to be more dynamic, including two abundant 3’ truncations of passenger strands in wildtype animals (Fig. S7C-D, Tables S10 and S11). Interestingly, passenger strand isomiRs were reduced in miR-2 mutants compared to wildtype, suggesting increased precision of generating the expected mature sequence (Fig. S7C-D). The miR-2-M2 and miR-2-M3 mutants, which asymmetrically select the miR-2 passenger strand (Fig. 3F), also altered the isomiR frequency of the miR-2 guide strand (Fig. S7C-D). Thus, while most of the miRNA mutants do not significantly affect isomiR populations, changes in isomiR populations were more frequently observed for passenger strands of highly asymmetric miRNAs. It is possible that certain isomiRs have altered duplex properties that promote their own strand selection, thereby leading to their increased representation relative to the corresponding canonical miRNA, without impacting the actual rate of isomiR formation. For example, in miR-2-M2 mutants, the most abundant guide strand isomiR was a truncation of the 5’ cytosine resulting in a more favorable 5’ adenine (Fig.S7C-D). Collectively, our data support a model where the identity and base-pairing status of 5’ nucleotides promotes miRNA strand selection but is not sufficient to explain strand preference in vivo.

### Identification of duplex asymmetries associated with miRNA strand choice

As we observed asymmetric strand selection for duplexes with symmetrical 5’ ends (Figs. 2 and 3), we hypothesized that additional duplex features influence miRNA asymmetry in vivo. As we were unable to identify additional asymmetries within the first four nucleotides on the 5’ end of guide and passenger strands (Fig. 4A), we next turned our attention to the 3’ end of duplexes and identified two nucleotide biases (Fig. 4B). The first bias was an adenine in the third-to-final position of miRNA passenger strands, which was noticeably enriched among the 30 highest expressed miRNAs in L4-staged animals (Fig. 4B) that represent more than 90% of total miRNA reads and have more confident passenger strand annotations (*37*) (Fig. 4C). As miRNA duplexes typically contain single stranded 2-nucleotide 3’ overhangs (*38, 39*), this adenine bias likely reflects base paring to the preferred 5’ uracil of guide strands. The second bias was the presence of a 3’ terminal cytosine on miRNA passenger strands, which was less frequently observed on guide strands and nearly absent from the 3’ ends of highly expressed miRNAs (Fig. 4B). As the levels of many miRNAs are developmentally regulated, we next examined whether this 3’ cytosine bias was present throughout animal development or specific to the L4-stage. Using an existing small RNA sequencing dataset (*29*), we quantified the observed 3’ nucleotide identity of miRNA guide and passenger strands throughout *C. elegans* development (Fig. 4D). Throughout animal development, we found that very few guide strands ended with a 3’ cytosine, whereas cytosine was among the most frequent 3’ nucleotides of passenger strands (Fig. 4D). Thus, 3’ nucleotide asymmetry is associated with miRNA strand choice throughout *C. elegans* development.

**Fig 4.**
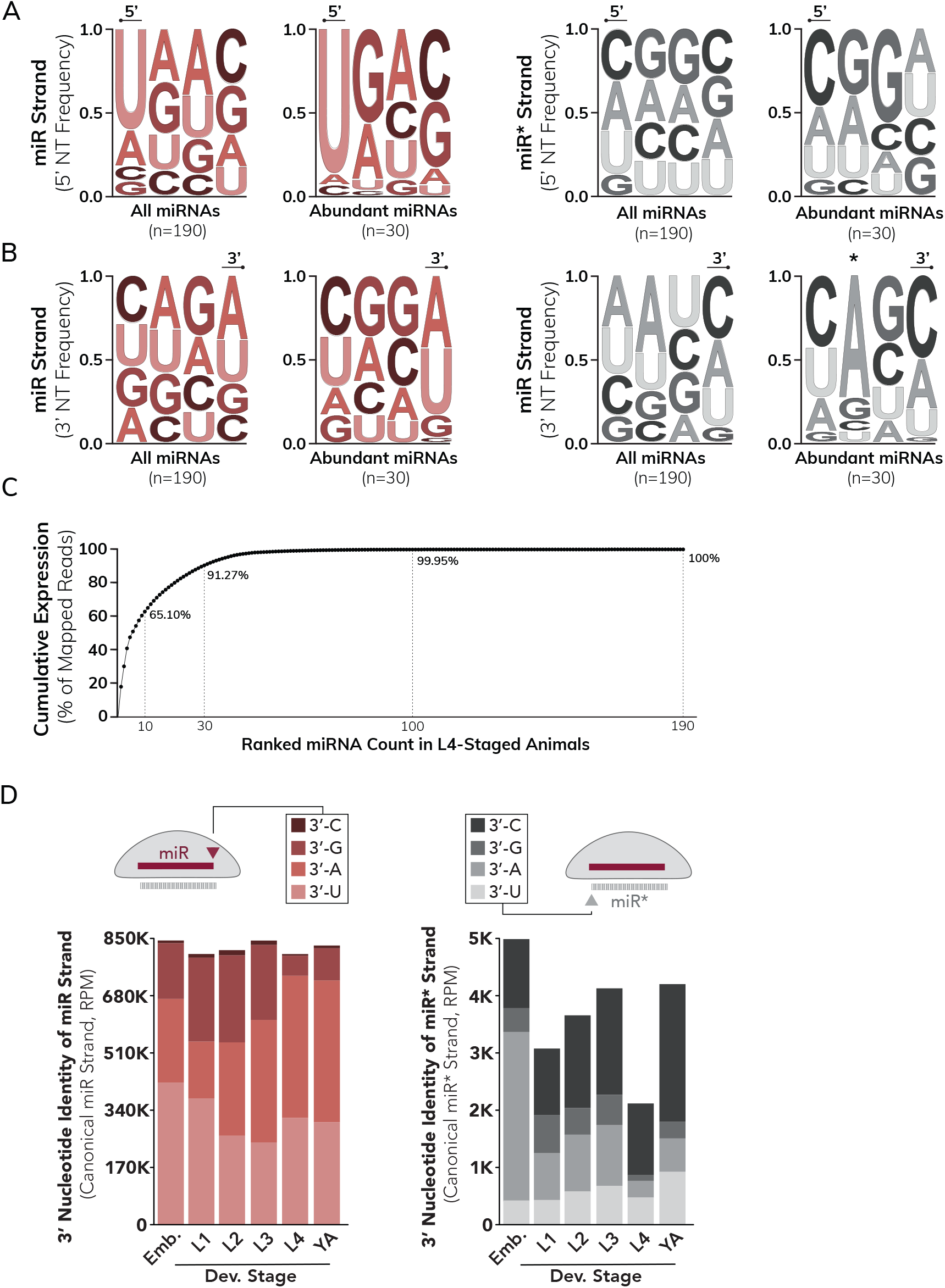
*C.elegans* miRNA duplex asymmetries. (A) Sequence logo representation of 5’ nucleotide frequencies for *C. elegans* miRNA guide (left) and passenger (right) strands. (B) Sequence logo representation of 3’ nucleotide frequencies for *C. elegans* miRNA guide (left) and passenger (right) strands. Asterisk indicates the third nucleotide from the 3’ duplex end of the passenger strand that is adjacent to the 3’ dinucleotide overhang and is expected to base pair to the 5’ nucleotide of the guide strand. (A-B) Sequences are given in 5’-3’ orientation, with the 5’ or 3’ terminal nucleotides indicated by a bulbed line above. Sequence logos were created using the weblogo3 server and colors were modified using Adobe illustrator. (C) Cumulative expression of C. elegans miRNAs at L4-stage. Each dot represents one miRNA (guide and passenger strands), plotted in order of most abundant to least abundant. The 10 most abundant miRNAs account for ∼65% and the 30 most abundant miRNAs represent >90% of all miRNA sequencing reads at the L4 stage. (D) Quantification of 3’ nucleotide frequency for miRNA guide (left) and passenger (right) strands throughout *C. elegans* development. Stacked bar graphs represent the normalized abundance of canonical miRNA guide or passenger strands (not including isomiRs) sorted by terminal 3’ nucleotide identity. Schematic (top) illustrates an example of a miRNA duplex loaded into Argonaute (gray) and the arrowhead indicates the position of the 3’ nucleotide of the guide (left) or passenger (right) strand. Developmental small RNA sequencing data were acquired from a previous study *(29)*.

### A 3’ cytosine on miRNA passenger strands promotes strand selection in *C. elegans*

To examine whether 3’ nucleotide identity influences miRNA strand selection in vivo, we created an additional set of *mir-58* mutations that alter 3’ nucleotide identity in the wildtype or symmetrical miR-58-M6 duplexes (Fig. 5, Table S2). The passenger strand of miR-58 has a 3’ cytosine and the guide strand has a 3’ uracil (Fig. 5A). We generated mutants where the 3’ nucleotide of both strands were swapped, or the 3’ nucleotide of one strand was mutated to produce duplexes containing symmetrical 3’ nucleotides (Fig. 5A). While swapping the 3’ nucleotide of the wild type miR-58 duplex in miR-58-M14 mutants was not sufficient to reverse strand selection, swapping the 3’ nucleotides of the miR-58-M6 duplex in miR-58-M16 mutants reversed strand preference (Figure 5A, Tables S5 and S6). This observation suggests a novel role for 3’ nucleotide identity in directing miRNA strand choice. Interestingly, in the miR-58-M6 background, we observed near equal levels of both guide and passenger strands for the miR-58-M17 mutant where both strands have 3’ cytosine, but asymmetric selection of the miR-58 guide strand in miR-58-M18 mutants where both strands have 3’ uracil (Fig. 5A). Thus, a specific preference for 3’ cytosine on passenger strands appears to regulate strand selection of miR-58.

**Fig 5.**
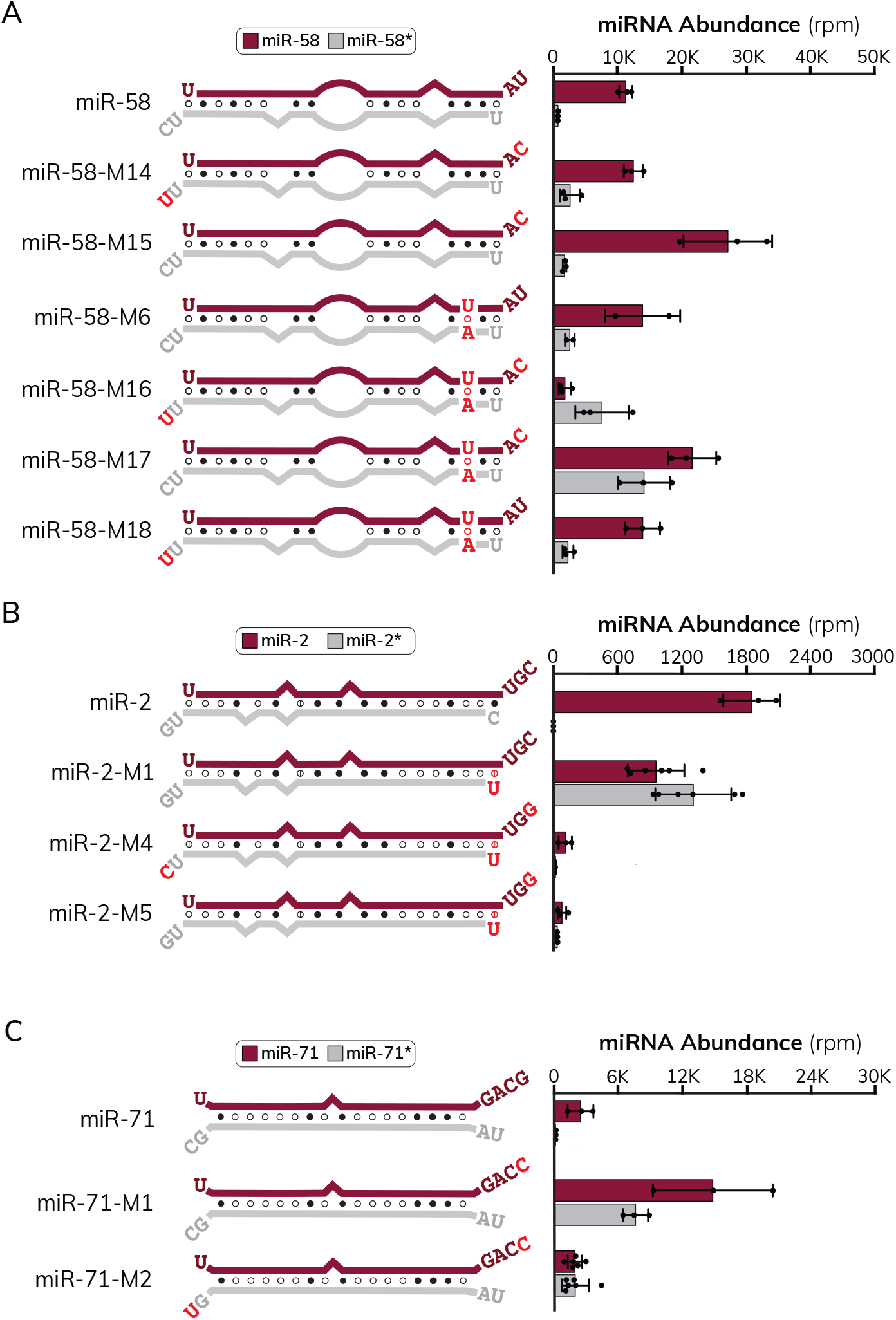
A 3’ cytosine on miRNA passenger strands is favorable for guide strand choice in *C. elegans*. (Left) Schematic representations of (A) miR-58, (B) miR-2 and (C) miR-71 duplex mutations that affect 3’ nucleotide identity. (A-C) The guide strand is shown in maroon, and the passenger strand is shown in gray. The nucleotide composition of dinucleotide overhang are shown for each strand. Note that the guide strand of miR-2 has an atypical trinucleotide overhang at its 3’ end. Mutated nucleotides are indicated by red text. U:A pairs are illustrated by an open circle, G:U wobble pairs are indicated by an open circle with a vertical line and G:C pairs are represented by closed circles. RNAcofold was used to predict duplex structures at 20°C. (Right) Quantification of small RNA sequencing results for (A) miR-58 (B) miR-2, and (C) miR-71. Data are presented as mean ± standard deviation, with each dot representing quantification from a single library using the NEXTFLEX v4 Small RNA Sequencing Kit. Maroon bars correspond to miRNA guide strands and gray bars represent miRNA passenger strands. Normalization was performed by dividing raw read counts by total mapped miRNA reads and multiplying by one million to acquire reads per million (rpm) values.

We next examined 3’ nucleotide mutations of *mir-2*, which has a 3’ guanine on the passenger strand and a 3’ cytosine on the guide strand (Fig. 5B). To minimize the effect of 5’ nucleotide identity, we mutated the miR-2-M1 duplex where both miR-2 strands have a symmetrical 5’ uracil (Fig. 5B, Table S1). The miR-2-M4 mutation switches the 3’ nucleotides of each miR-2-M1 strand and the miR-2-M5 mutations mutates the 3’ nucleotide of the miR-2-M1 passenger strand so that both strands have a symmetrical 3’ guanine (Fig. 5B). The miR-2-M1 mutation reversed strand preference of miR-2 (Figs. 2B and 5B) presumably due to the 3’ cytosine on the canonical miR-2 guide strand (passenger strand of miR-2-M1) (Fig. 5B). This passenger strand accumulation in miR-2-M1 was relieved by mutation of the 3’ cytosine on the miR-2 guide strand in miR-2-M4 and miR-2-M5 mutants, consistent with a 3’ cytosine directing strand selection of miR-2 (Fig. 5B). Analysis of miR-2 (Fig. S7C-D, Tables S10 and S11) and miR-58 isomiRs (Fig. S6, Tables S7 and S8) indicated that the 3’ nucleotide variants were processed correctly. Furthermore, introducing a 3’ cytosine to the guide strand of miR-71 (Fig. 5C, Table S2) promoted selection of the miR-71 passenger strand without affecting isomiR populations (Fig. S8, Table S12). Introducing a 3’ cytosine to either strand of the miR-48 duplex (Table S2) led to a small accumulation of the opposing strand without affecting miR-48 processing (Fig. S8, Table S13). Collectively, these data show that 3’ nucleotide identity influences the strand selection of a wide range of miRNAs in vivo.

The nucleotide composition of 3’ overhangs may influence strand asymmetry, as we observed modest differences in strand selection when we swapped 3’ nucleotides or the dinucleotide overhangs of miR-58 duplexes (Fig. S9). However, we did not identify obvious nucleotide asymmetries beyond the terminal nucleotide of 3’ ends, so we chose to focus on the role of 3’ nucleotide identity in this study (Fig. 4B). We next asked whether 3’ nucleotide identity might be directly recognized by Argonaute. To address this, we mutated a putative 3’ nucleotide binding pocket (*17*) that is positioned near the 5’ nucleotide binding pocket in the MID domain (*13, 14, 25*) of the *C. elegans* miRNA-associated Argonaute ALG-1 (Fig. S10A-B). Although small RNA sequencing did not identify miRNA strand switching in ALG-1^7A^ mutants, we observed accumulation of several miRNA passenger strands (Fig. S10C, Table S6), suggesting that the interaction between 3’ nucleotides and Argonaute may, at least in part, promote accurate strand selection. Collectively, our results suggest that a 3’ cytosine on miRNA passenger strands promotes guide strand selection and is sufficient to direct strand selection of miRNAs with symmetrical 5’ duplex ends.

### A 3’ nucleotide identity plays a conserved role in directing miRNA choice

We next asked whether 3’ nucleotide identity influences miRNA strand selection in an evolutionarily conserved fashion. We used the shERWOOD UltramiR system (*40*) to introduce a set of miR-58 variants into human HEK293T cells (Fig. 6A). The human genome does not encode a miR-58 homolog, which allowed us to track the strand preference of the exogenous miR-58 variants. We first transfected the wildtype miR-58 and symmetrical miR-58-M6 duplexes as well as their respective 3’ nucleotide swapped variants miR-58-M14 and miR-58-M16 (Fig. 6A). Our small RNA sequencing (Table S14) showed that the canonical miR-58 guide strand is asymmetrically selected, like what is observed in *C. elegans* (Fig. 6A). However, unlike in *C. elegans*, both strands of the symmetrical miR-58-M6 duplex appear to be selected at near equal frequency (Fig. 6A), suggesting an inherent difference in miRNA strand selection between *C. elegans* and humans. Furthermore, we found that passenger strand levels were reduced in the 3’ nucleotide swapped variants miR-58-M14 and miR-58-M16 that each have a 3’ cytosine on the miRNA guide strand (Fig. 6A), suggesting 3’ cytosine is less favored in human cells. Each of the miR-58 variants appeared to be processed correctly based on isomiR analysis (Fig. S11, Table S15). However, an additional set of mutations that we designed in a modified version of the human miR-339 duplex were not processed as expected (Fig. S11, Table S15), despite a modest effect on strand asymmetry by those mutations (Table S14).

**Figure 6.**
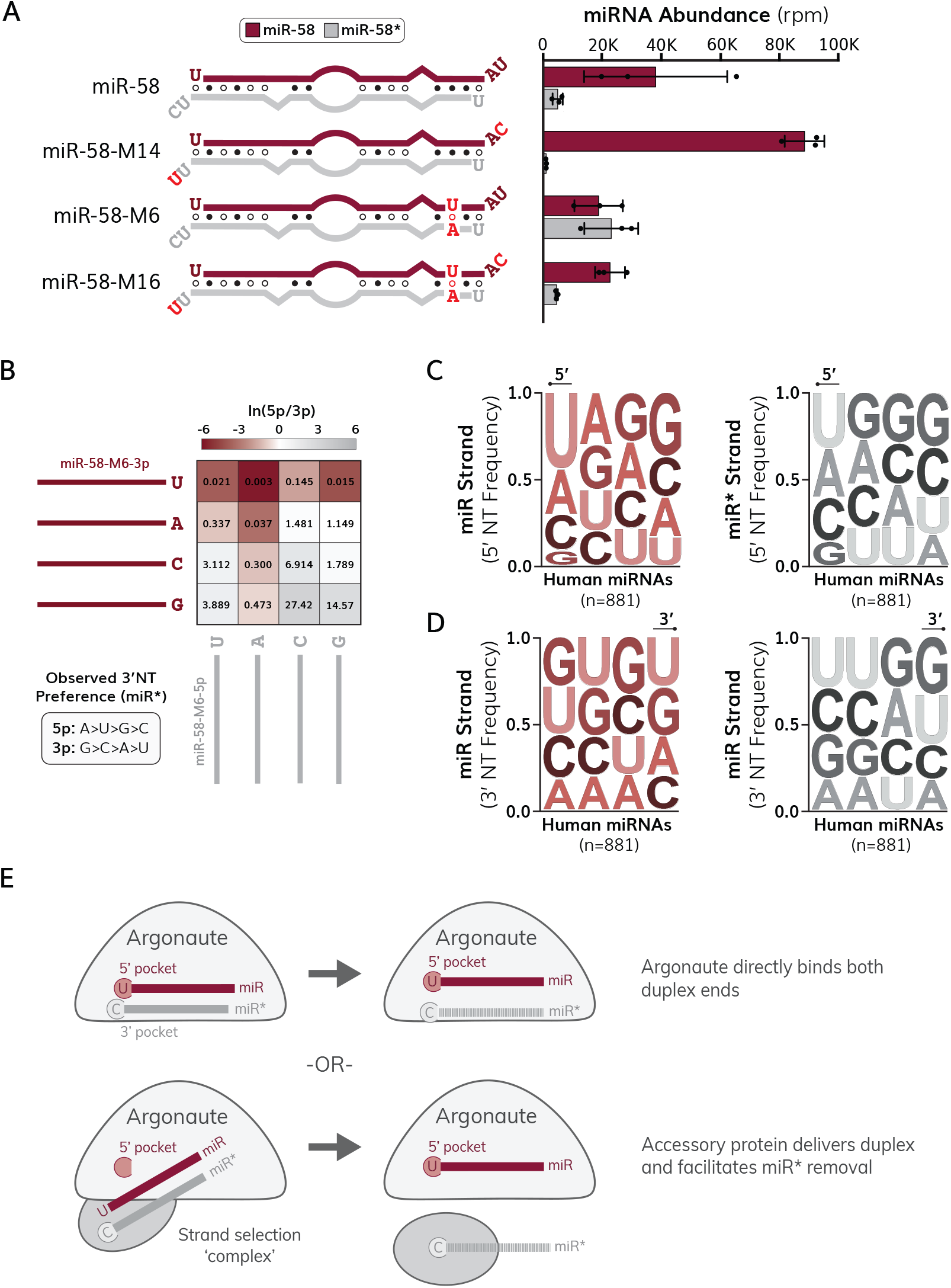
3’ nucleotide identity influences miRNA strand asymmetry in human HEK293T cells. (A) (Left) Schematic representations of exogenous miR-58 duplexes transfected into HEK293T cells that alter 3’ nucleotide identity. The canonical miR-58 guide strand (from *C. elegans*) is shown in maroon, and the passenger strand is shown in gray. The nucleotide composition of dinucleotide overhang are shown for each strand. U:A pairs are illustrated by an open circle, G:U wobble pairs are indicated by an open circle with a vertical line and G:C pairs are represented by closed circles. RNAcofold was used to predict duplex structures at 37°C. (Right) Quantification of small RNA sequencing results for transfected miR-58 variants. Data are presented as mean ± standard deviation, with each dot representing quantification from a single library using the NEXTFLEX v4 Small RNA Sequencing Kit. Maroon bars correspond to canonical miRNA guide strands and gray bars represent canonical miRNA passenger strands. Normalization was performed by dividing raw read counts by total mapped miRNA reads and multiplying by one million to acquire reads per million (rpm) values. (B) Absolute quantification of miR-58-M6 duplex variants comprising each combination of 3’ nucleotides on each duplex strand. Heatmap illustrates calculated ln(5p/3p) strand ratios where a deeper maroon color indicates preference for the miR-58-M6-3p (guide) strand. The non-transformed 5p/3p strand ratios are given in each box. (C) Sequence logo representation of 5’ nucleotide frequencies for human miRNA guide (left) and passenger (right) strands. (D) Sequence logo representation of 3’ nucleotide frequencies for human miRNA guide (left) and passenger (right) strands. (C-D) Sequences are given in 5’-3’ orientation, with the 5’ or 3’ terminal nucleotides indicated by a bulbed line above. Sequence logos were created using the weblogo3 server and colors were modified using Adobe illustrator. (E) Hypothetical models of how 3’ nucleotide identity influences miRNA strand selection. (Top) The MID domain of Argonaute may directly interact with the 5’ nucleotide of the guide strand and the 3’ nucleotide of the passenger strand. Such interaction may induce a conformational change that promotes duplex wedging and release of the passenger strand. (Bottom) Alternatively, additional factors may interact with miRNA 3’ nucleotides to form a strand selection complex. Dissolution of such complex could facilitate guide strand retention within Argonaute and passenger strand removal.

To characterize the 3’ nucleotide preference in human cells, we created several miR-58-M6 duplexes comprising 16 possible 3’ nucleotide combinations for each duplex strand (Fig. 6B). To quantify the levels of each miR-58 variant, we performed absolute quantification for each duplex strand (Fig. 6C, Table S16). We found that a 3’ guanine on the guide (3p) strand of miR-58-M6 and a 3’ uracil on the passenger (5p) strand of miR-58-M6 were most favorable for guide strand selection (Fig. 6B). Notably, when each miR-58-M6 strand contained a 3’ uracil or adenine, the 3p strand was highly favored whereas when each strand contained a 3’ guanine or cytosine the 5p strand was asymmetrically selected (Fig. 6B). These observations suggest that precursor origin may influence miRNA strand choice, or that additional duplex features, such as 3’ overhang composition, may play a prominent role in directing miRNA strand choice in human cells. We were unable to identify an obvious nucleotide bias at the 3’ end of human miRNA duplexes (Fig. 6C-D, Table S17), consistent with the idea that additional duplex features may influence strand asymmetry in human cells. Collectively, our results indicate that 3’ nucleotide asymmetry plays an evolutionarily conserved role in regulating miRNA strand choice. While it is unclear how 3’ nucleotide asymmetry is recognized, we propose two alternative hypotheses that are not mutually exclusive, where the 3’ nucleotide is directly bound by Argonaute or by an accessory protein to facilitate guide strand selection and passenger strand removal (Fig. 6E).

## Discussion

How does 3’ nucleotide identity promote miRNA strand choice? One possibility is that Argonaute directly engages both the 5’ nucleotide of the guide strand (*13, 14, 25*) and the 3’ nucleotide of the passenger strand to initiate duplex unwinding (Fig. 6E). Consistent with this idea, we found that mutating a putative 3’ nucleotide binding pocket on Argonaute (*17*) led to an accumulation of some miRNA passenger strands (Fig. S10C). Duplex binding may induce a conformational change of Argonaute, promoting passive wedging by its N-terminal domain (*41*) and coupling nucleotide recognition to duplex unwinding. Notably, mutations in *C. elegans* ALG-1 and its human homolog Ago2 affect miRNA strand asymmetry (*42, 43*), although it is currently unclear whether those mutations are connected to recognition of 3’ nucleotide identity.

An alternative, but not mutually exclusive, hypothesis is that Argonaute cooperates with additional factors that recognize 3’ nucleotide identity and bias strand selection (Fig. 6E). For example, the Dicer-interacting proteins TRBP and PACT regulate miRNA strand selection in humans (*44*–*46*), and Dicer itself can reposition the miRNA duplex after dicing (*45*), raising the possibility that Dicer hands off miRNA duplexes to Argonaute in a favorable orientation. Interestingly, the *C. elegans* genome lacks obvious homologs for both TRPB and PACT, which may at least partially account for the differences in strand selection and 3’ nucleotide preferences we observed between *C. elegans* and human cells. More broadly, variation in the protein composition of strand selection complexes could influence miRNA sorting by directing duplexes to distinct Argonaute proteins with different 3’ nucleotide preferences. Although 3’ nucleotide identity appears to play a conserved role in miRNA strand selection, it is worth noting that human miRNAs, unlike *C. elegans* miRNAs, lack an obvious global 3’ nucleotide asymmetry. This observation highlights the value of model systems such as *C. elegans* in understanding the basic mechanisms of miRNA strand selection.

In addition to identifying a novel determinant of miRNA strand choice, our findings suggest that 3′ nucleotide identity may contribute to the dynamic regulation of miRNA strand asymmetry. While both regulation and dysregulation of miRNA strand asymmetry have previously been reported (see review (*18*)), current models of miRNA strand selection lack a dynamic component to explain context-dependent changes in strand preference (*17*). As 3’ isomiRs are typically more frequent than 5’ isomiRs (*29, 47, 48*), changes in 3’ end identity reported in developmental (*29, 49*–*51*), cellular (*52, 53*) or disease (*54*) contexts may underly miRNA strand dynamicity. Some 3’ modifications, such as miRNA tailing and trimming (*36, 55*), likely occur after strand selection and are therefore unlikely to affect miRNA strand choice. Other 3’ modifications, such as adenylation or uridylation, often target pre-miRNA molecules (*48, 56*–*58*) and are more likely to influence strand asymmetry. Consistently, uridylation of human miR-324 leads to altered strand selection, although uridylation of miR-324 may primarily influence Dicer-dependent processing (*59*). Intriguingly, cytidylation of miRNA precursors has been observed in Arabidopsis (*60*) and the miR-83-3p guide strand is cytidylated in *C. elegans* (*55*). As our data indicate that a 3’ cytosine on miRNA passenger strands is favorable for strand selection, cytidylation of one strand could bias loading toward the other duplex strand. As these passenger strand modifications would further favor guide strand selection, they are likely to be underrepresented in small RNA sequencing datasets, potentially explaining why such modifications have not yet been identified. Additional work will be needed to fully understand how 3’ nucleotides are recognized and how changes in 3’ nucleotide identity may regulate miRNA strand selection across different contexts and disrupt miRNA strand selection in human disease.

## Materials and Methods

### *C. elegans* Strains and Genetic Analysis

A complete list of *C. elegans* strains used in this study is provided in Table S18. All strains were derived from the wild-type N2 genetic background. Strains were maintained at 20°C under standard growth conditions (*61*). Nematode growth medium (NGM) plates that were seeded with *Escherichia coli* OP50 were used as a food source. For developmental stage-specific analysis, animals were synchronized using standard alkaline hypochlorite treatment (*62, 63*) and harvested when most (>90%) of the larval population reached the desired developmental stage. For body length measurements, adult animals were imaged on a Leica DM6 upright microscope. Adobe illustrator was used to quantify body length by measuring the path length of a curved line drawn through the center of the animal from the head to base of the tail. For drug assays, young adult worms were added to plates containing 200 µM levamisole (*31*) or 1mM aldicarb (*32*) and scored for paralysis at indicated time intervals. Paralysis was defined as the inability for worms to move in response to a touch stimulus from a platinum wire.

### Profiling miRNA Duplex Characteristics

Mature *C. elegans* and human miRNA sequences were retrieved from miRbase v 22.1 (*37*). Guide and passenger strand annotations were determined based on which strand had the most reported reads on miRbase. 5’ nucleotide information was compiled for 5p and 3p strands. RNAcofold from the ViennaRNA Package 2.0 (*64*) was used for duplex folding and to calculate thermodynamic end stability at 20°C as previously described (*28*). Predictions of miRNA strand selection were calculated using twin-drive model as previously described (*17*).

### CRISPR/Cas9 Genome Editing

*Streptococcus pyogenes* Cas9 (IDT, Alt-R® S.p. Cas9 Nuclease V3), single-stranded repair templates (IDT, custom ultramers), crRNA and tracrRNA were commercially sourced. A full list of oligonucleotides used in this study is provided in Table S19. Microinjection and screening were performed using *dpy-10* co-conversion as previously described (*65*). Cas9 was injected at a final concentration of 2.65 µM and all injection mixes included 200 µM KCl and 7.5 µM HEPES [pH 7.4]. All crRNAs were injected at a final concentration of 50 µM, except *dpy-10* crRNA was injected at a final concentration of 5 µM. Equimolar tracrRNA was included in all injection mixes. Single stranded DNA oligonucleotides were used to facilitate homology-directed repair and were included at final concentrations of 6 µM or 3 µM (*dpy-10*).

### RNA Extraction and Isolation

Immediately after sample collection, samples were centrifuged and flash-frozen in liquid nitrogen and stored at -80°C until ready for use. Pellets were thawed in 250 µL RNase-free water (final volume). 1 mL of Trizol (Invitrogen) was added and the mixture was continuously vortexed for 10 minutes. Afterwards, 212 µL of chloroform was added and vortexed for an additional minute. The aqueous and organic layers were separated by centrifugation at 19000 x g for 10 minutes at 4°C. The aqueous phase was then recovered and re-extracted using equal volume of phenol:chloroform:isoamyl alcohol (25:24:1, VWR). The RNA was precipitated for 2 hours at -20°C using an equal volume of isopropanol. 1.8 µL of Glycoblue (Invitrogen) was added as a co-precipitant. The RNA pellets were washed using 80% (v/v) ethanol and allowed to air-dry on ice for 10 minutes. The dried pellets were dissolved in RNase-free water and stored at -80°C. RNA concentration was determined using a Nanodrop spectrophotometer.

### Small RNA Sequencing and Analysis

For preparation of small RNA libraries, total RNA was first size selected using a 15% denaturing gel using UreaGel (National Diagnostics) reagents. Single-stranded 18mer (AGCTGCCAGCAACTCTAA) and 26mer (TAGTATGTAGCTGCCAGCAACTCTAA) were combined at a final concentration of 5 µM and used as a molecular ladder in an empty lane. Gels were soaked in 1XTBE solution containing 1X SYBR Gold stain (Invitrogen) for 10 minutes and subsequently visualized under UV light. Small RNAs migrating between the 16mer and 26mer markers were excised from the gel and soaked in 750 µL of TE buffer supplemented with 0.3 M NaCl overnight while shaking at 550 rpm at 7°C. The solution was transferred to a 0.45 µm column and centrifuged at 4°C for 10 minutes at 2500 x g. Small RNAs were precipitated for 2 hours at -20°C using equal volume of isopropanol and 10% sample volume 3M NaOAc. 1.5 µL of Glycoblue (Invitrogen) was used as a co-precipitant. The small RNA pellets were washed once using 80% (v/v) ethanol and allowed to air-dry on ice for 10 minutes. The dried pellets were dissolved in 11 µL of RNase-free water. Size-selected small RNA samples were used to make libraries using the Nextflex Small RNA-Seq Kit v3 (PerkinElmer) or the Nextflex Small RNA-Seq Kit v4 (Revvity) according to manufacturer instructions. Please note that the Nextflex Small RNA-Seq Kit v3 (PerkinElmer) was discontinued during this study. We separated small RNA sequencing data acquired from the v3 (Table S5) and v4 kits (Table S6) and only made direct comparisons for samples that were sequenced using the same kit, as we observed differences in normalized read counts for several miRNAs when using different kits. All small RNA libraries were sequenced at the Kansas State University Integrated Genomics Facility using an Illumina NextSeq 500 instrument at 75 cycles.

Small RNA sequence data were processed and analyzed as previously described (*29*) The cutadapt tool (*66*) was used to process and trim the sequencing reads. The 3’ adapter sequences were first removed using the parameters “-a ATCTCGTATGCCGTCTTCTGCTTGX” and demultiplexed using the parameters “e 1 -a file$:barcodes.fasta”, where the “barcodes.fasta” contains a list of barcode sequences in fasta format. The 3’ primer sequence was then removed from each read using the parameters “-a TGGAATTCTCGGGTGCCAAGGAACTCCAGTCAC”. For the Nextflex Small RNA-Seq Kit v3, each end of the processed sequencing reads contained four-nucleotide randomers that were removed using cutadapt. The final four nucleotides of each sequencing read were trimmed using the parameters “-u -4”. The first four nucleotides were trimmed using the parameters “-u 4 -m17 -M29”, which also removed processed reads that were shorter than 17 or longer than 19 nucleotides. Read quality was examined using the fastqc tool (*67*). Reads were aligned to a list of mature *C. elegans* miRNAs using bowtie (*68*). The following command was used for bowtie alignments: “-x miR -a--best --strata --norc -v 3”, where “miR” is a bowtie build assembled using a fasta file containing all *C. elegans* mature miRNA sequences obtained from miRbase. The samtools tool (*69*) was used to generate, sort and index bam files to generate read count data. A custom bash script was used to identify and quantify isomiR abundance from fastq files. Normalization (reads per million) was performed by multiplying the number of mapped reads for individual miRNA loci by the total number of mapped miRNA reads divided by one million.

### qPCR and Absolute Quantification

The QuantiNova SYBR Green RT-PCR kit (Qiagen) was used for standard qPCR reactions according to manufacturer instructions. Final reaction volumes were adjusted to 5 µL and 100 ng of total *C. elegans* RNA was used as a template. Absolute quantification of miRNAs using Taqman was performed as previously described (*70*). Standard curves were first made using known quantities (0.6ng/µL, 0.6×10^-2^ ng/µL, 0.6×10^-4^ ng/µL, 0.6×10^-6^ ng/µL, and 0.6×10^-8^ ng/µL) of commercially purchased (IDT) single-stranded RNA oligonucleotides prepared from a 2 µg/µL stock. All reactions were carried out at a final volume of 10 µL using the Taqman™ MicroRNA Reverse Transcription Kit (Thermofisher) and Taqman™ Universal Master Mix II, no UNG (Thermofisher) according to manufacturer instructions. Custom or predesigned Taqman™ Small RNA Assays (ThermoFisher) were used at 1X final concentration. 1 ng of total *C. elegans* RNA was used as a template for reverse transcription (10 µL reaction volume) and 0.67 µL of the cDNA product was used for experimental Taqman assays. The ABI 7500 Real-Time PCR System (Applied Biosystems) was used to run all qPCR and Taqman reactions. A full list of oligonucleotides and Taqman assays used in this study is provided in Table S19.

### UltramiR Plasmid Construction

A full list of plasmids used in this study is provided in Table S20. Each miRNA variant was cloned into the pLMN-ZsGreen-Neomycin-shRNA vector (transOMIC) and transformed into NEB^®^ Stable Competent *E. coli* (High Efficiency) according to manufacturer instructions. The pJM1 through pJM7 plasmids were created by Gibson assembly of Gblock gene fragments (IDT) (Table S19) and HpaI/MluI (NEB) digested pLMN-ZsGreen-Neomycin-shRNA vector using the Gibson Assembly^®^ Cloning Kit (NEB) according to manufacturer instructions. The pJM10 through pJM23 plasmids were created by performing site-directed mutagenesis of the pJM2 plasmid to mutate the nucleotides corresponding to the 3’ end of the 5p or 3p strands using the Q5^®^ Site-Directed Mutagenesis Kit (NEB) according to manufacturer instructions. The primers used for site-directed mutagenesis are listed in Table S19. All clones were verified by Sanger sequencing prior to transfection

### Human Cell Culture and Transfection

Human HEK 293T cells were seeded in 12-well plates containing 1 mL DMEM with 10% FBS and transfected when reaching ∼90% confluency. In one tube 1 µg of UltramiR plasmid was added to 100 µL of Opti-MEM™ (Thermofisher) and vortexed. In a second tube, 5 µL of 1 mg/mL polyethylenimine (PEI) reagent was added to 100 µL of Opti-MEM™ (Thermofisher), vortexed and incubated at room temperature for 5 min. The contents of the second tube were subsequently pipetted into the first tube, vortexed and incubated at room temperature for 15 min. The mixture was then carefully added to the cell media dropwise. After 6 hours, the cell media was changed with fresh warmed DMEM with 10% FBS. 48 hours post-transfection, cells were imaged for ZsGreen expression (Figure S12), harvested in 1 mL of Trizol (Invitrogen), frozen and stored at -80ºC until use.

### Statistical Analysis

R statistical software was used to calculate all statistics. All *P*-values were calculated using two-tailed *t*-tests assuming equal variance among samples. All statistics are presented as mean ± one standard deviation unless otherwise specified.

## Data Availability

Small RNA sequencing data files have been uploaded to the SRA database (PRJNA1378235) and will be released upon final publication. All strains and plasmids used in this study are available upon request. All other data supporting the findings of this study are available from the corresponding author on reasonable request.

## Acknowledgements

We thank members of the Zinovyeva lab for helpful discussions. We thank Hui Tian, Sarah Coffey and Katherine Adams (Hwang) for technical assistance in generating some worm strains, and Alina Akhunova from the Kansas State Integrated Genomics Facility for technical expertise in generating small RNA sequencing data. We are grateful to Anne Hart for sharing *mir-2* mutant strains. Some of the strains used in this study were provided by the *Caenorhabditis* Genetics Center (CGC) which is funded by NIH Office of Research Infrastructure Programs (P40 OD010440). This work was supported by grants from the National Institutes of Health (R35GM124828 to A.Z. and R01GM117406 to C.M.H.), the Kansas INBRE program (P20 GM103418) and the Johnson Cancer Center at Kansas State University. J.C.M was supported by a fellowship from the NIH (F32GM148040).

## Author Contributions

J.C.M and A.Z. conceptualized and designed the experiments. A.Z and C.M.H supervised the research and acquired funding. J.C.M, S.K, H.O, H.C, S.Z, W.S, J.S and M.B carried out the experiments. J.C.M and A.Z. performed data analysis and interpreted results. J.C.M and G.P performed the required bioinformatics. J.C.M wrote the original manuscript draft, and all other authors edited the manuscript.

