## Supplementary Materials for "3’ Nucleotide Asymmetry Directs miRNA Strand Selection"

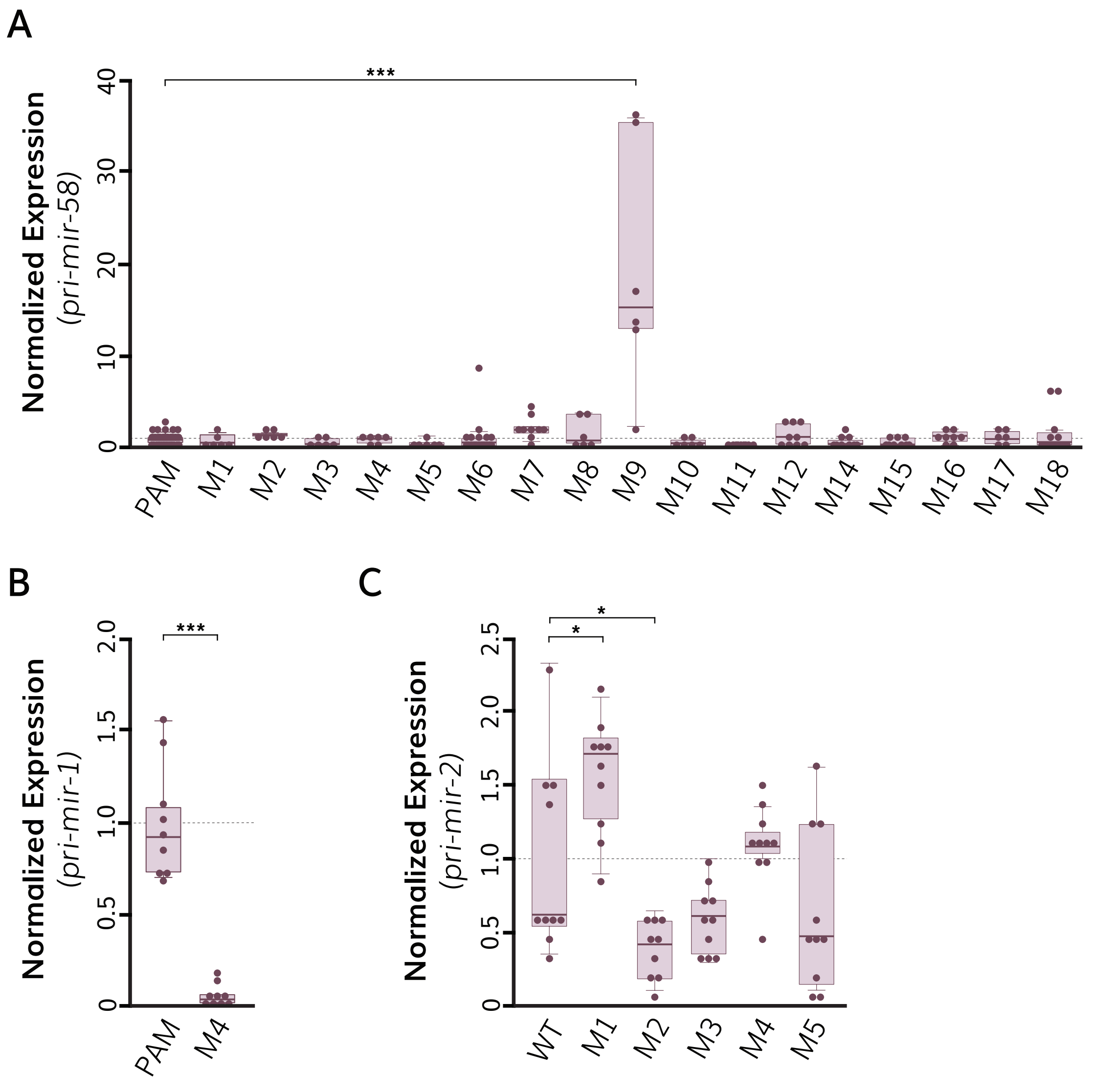


**Fig S1.** Quantification of (A) *pri-mir-58*, (B) *pri-mir-1* and (C) *pri-mir-2* levels in L4-staged miRNA mutants by qRT-PCR. (A-C) Each dot represents one qRT-PCR data point. Each sample was performed in duplicate and at least three independent replicates were included for each mutation. Data were first normalized to α-tubulin (*tba-1*) and subsequently normalized to appropriate PAM or wildtype (WT) controls. The dashed horizontal line indicates the self-normalized average for appropriate controls. Boxes range from the first to third quartile of the data. The thick bar represents the statistical median. Lines extend to the minimum and maximum data point excluding outliers that were defined as 1.5 times the interquartile range. Statistical analysis: *p<0.05, ***p<0.001.

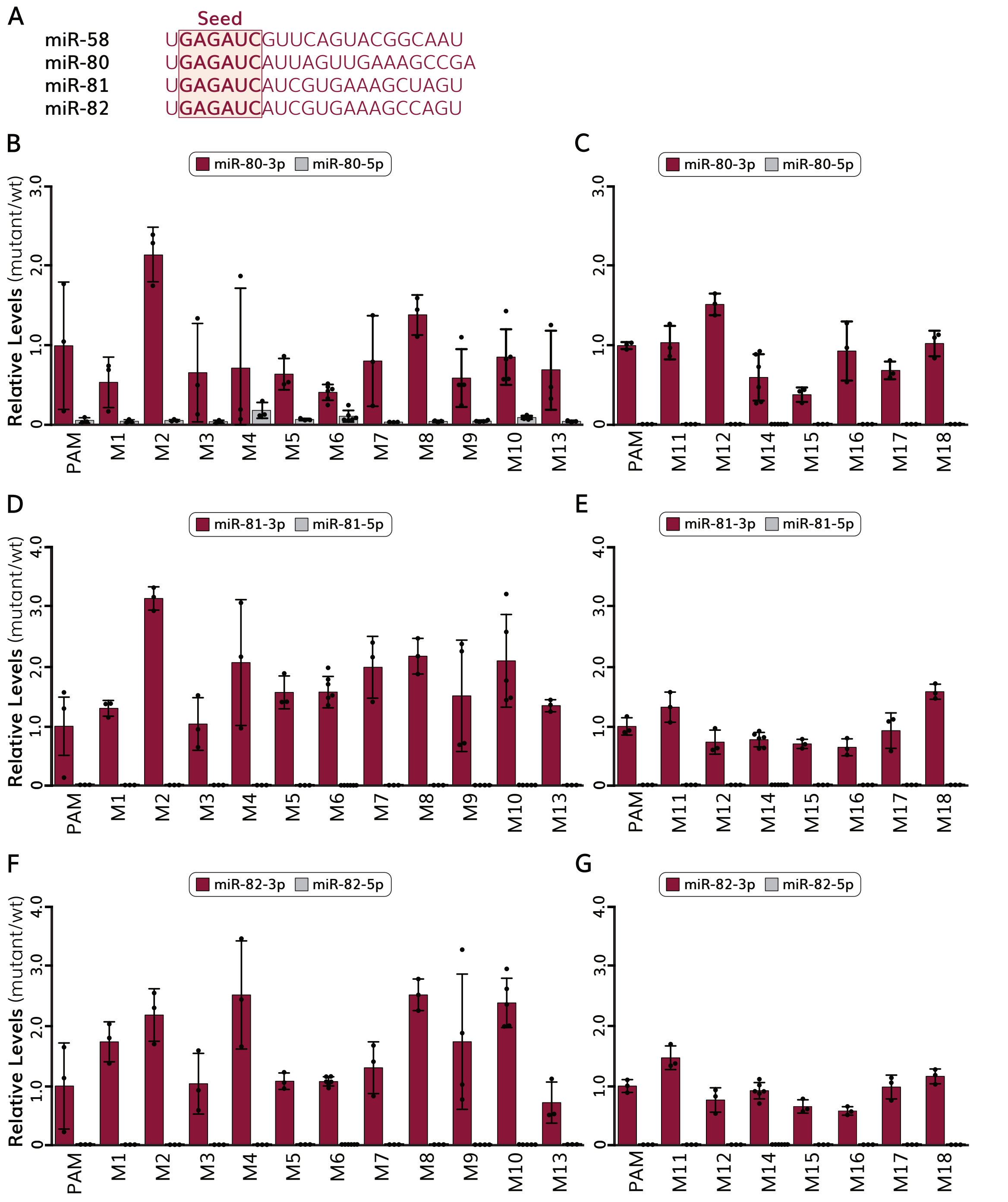


**Fig S2.** Analysis of miR-58 family miRNAs in miR-58 mutants. (A) Sequence alignment of the *C. elegans* miR-58 miRNA family members. (B-C) Quantification of mature miR-80 levels in L4-staged animals normalized to PAM controls. Libraries were created with the (B) Nextflex v3 small RNA sequencing kit or the (C) Nextflex v4 small RNA sequencing kit. (D-E) Quantification of mature miR-81 levels in L4-staged animals normalized to PAM controls. Libraries were created with the (D) Nextflex v3 small RNA sequencing kit or the (E) Nextflex v4 small RNA sequencing kit. (F-G) Quantification of mature miR-82 levels in L4-staged animals normalized to PAM controls. Libraries were created with the (F) Nextflex v3 small RNA sequencing kit or the (G) Nextflex v4 small RNA sequencing kit. (B-G) Data are presented as mean ± standard deviation, with each dot representing quantification from a single library. Maroon bars correspond to canonical miRNA guide strands and gray bars represent canonical miRNA passenger strands.

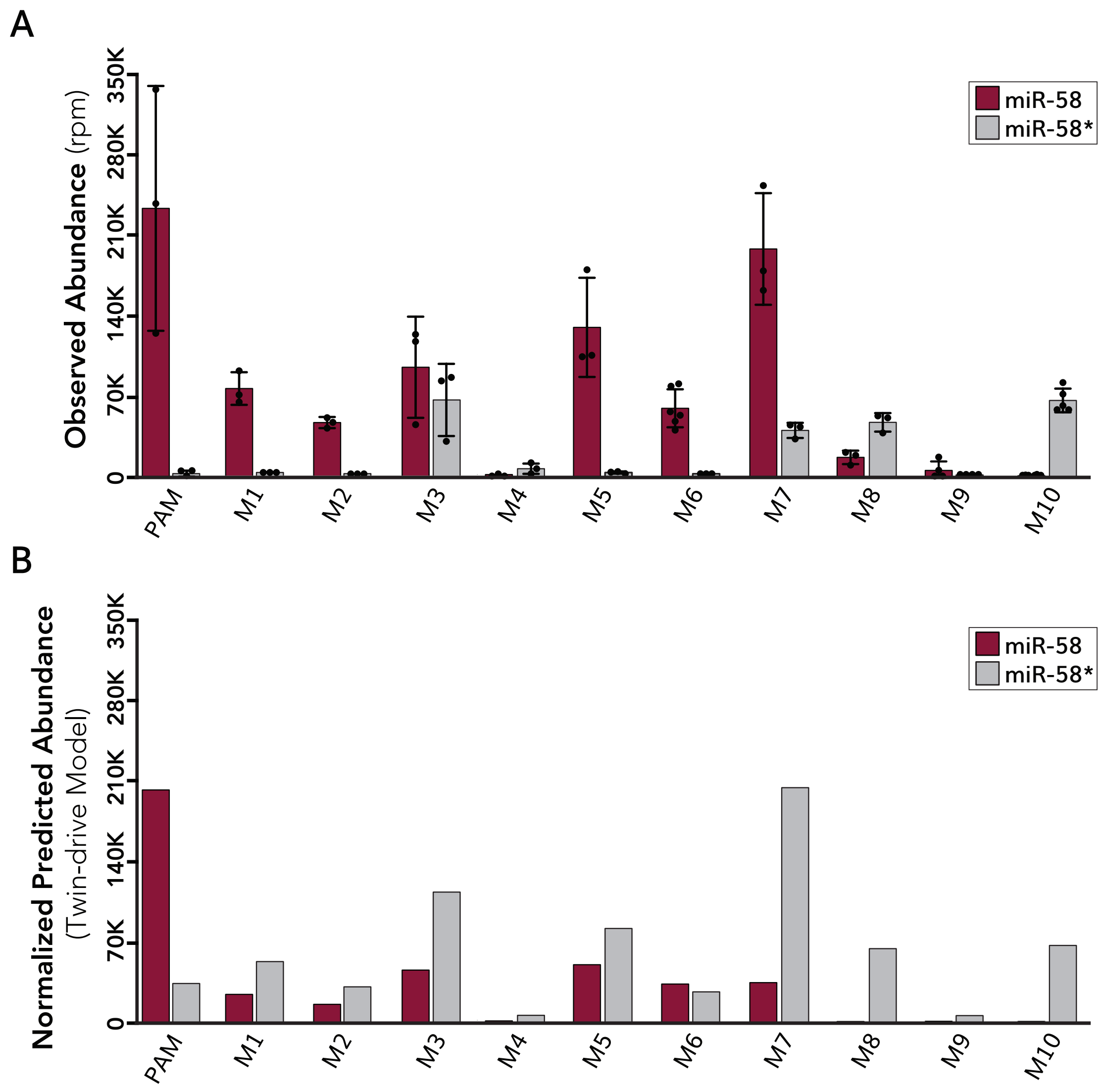


**Fig S3.** Comparison of observed miR-58 strand ratios to twin-drive model predictions. (A) Quantification of small RNA sequencing results for each miR-58 duplex strand. Data are presented as mean ± standard deviation, with each dot representing quantification from a single library using the NEXTFLEX v3 Small RNA Sequencing Kit. Maroon bars correspond to miRNA guide strands and gray bars represent miRNA passenger strands. Normalization was performed by dividing raw read counts by total mapped miRNA reads and multiplying by one million to acquire reads per million (rpm) values. (B) Predicted miR-58 strand abundance using twin-drive model. 5p/3p strand ratios were normalized to the observed abundance from small RNA sequencing of each miR-58 variant. Maroon bars correspond to miRNA guide strands and gray bars represent miRNA passenger strands.

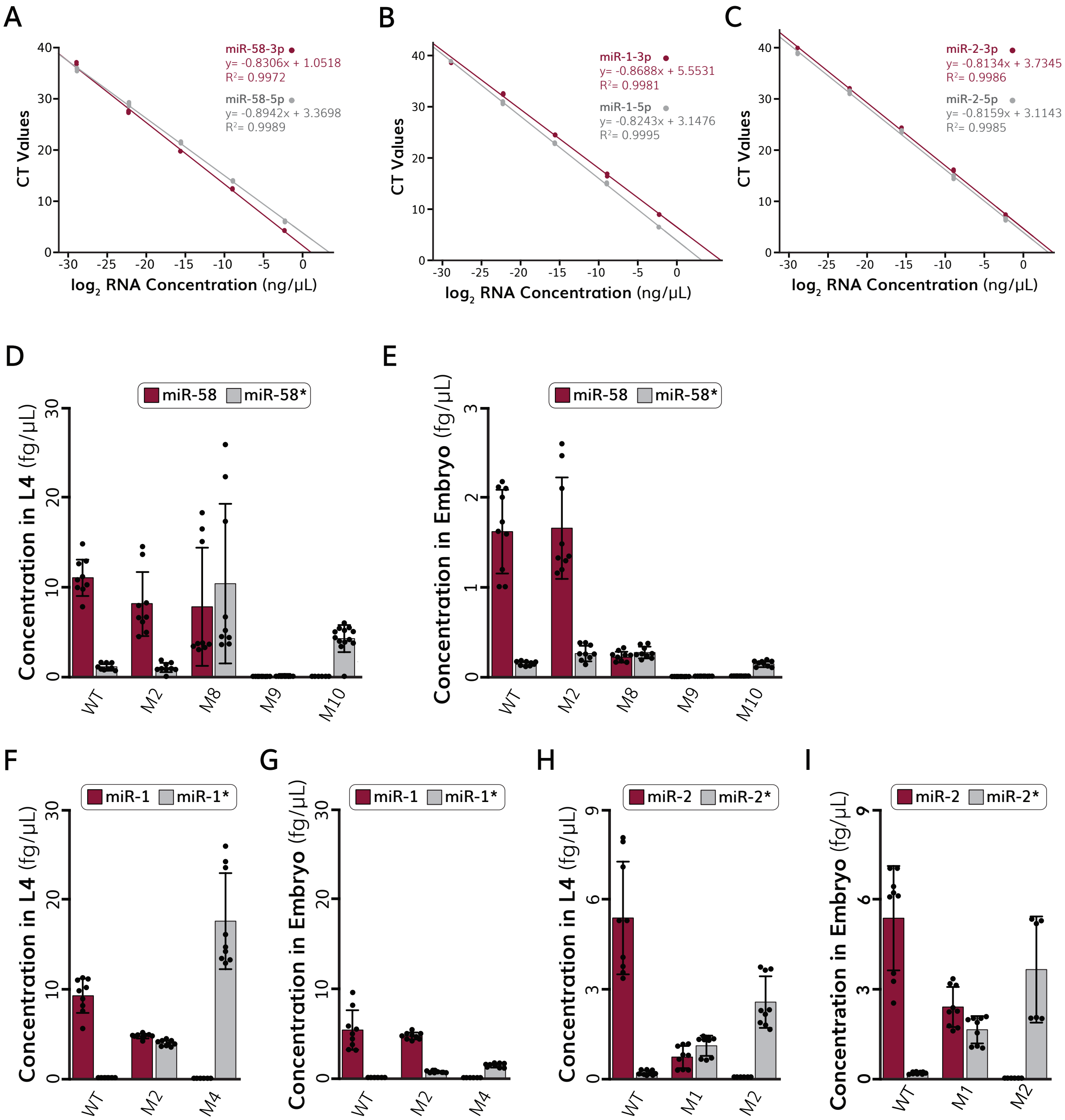


**Fig S4.** Absolute quantification of miRNA levels in representative miRNA mutants using miRNA Taqman assays. (A-C) Standard curves created using known amounts of guide and passenger strand RNA for (A) miR-58, (B) miR-1 and (C) miR-2. Linear regression and R2 coefficients are indicated for guide (maroon) and passenger (gray) strands. Each dot represents an independently prepared Taqman assay. (D-E) Absolute quantification of both miR-58 strands in (D) L4-staged or (E) embryonic miR-58 mutants. (F-G) Absolute quantification of both miR-1 strands in (F) L4-staged or (G) embryonic miR-1 mutants. (H-I) Absolute quantification of both miR-58 strands in (H) L4-staged or (I) embryonic miR-52 mutants. (D-I) Data are presented as mean ± standard deviation. Each sample was assayed in triplicate and at least three independent replicates were included for each mutation. Each dot represents quantification from a single Taqman assay. Maroon bars correspond to canonical miRNA guide strands and gray bars represent canonical miRNA passenger strands.

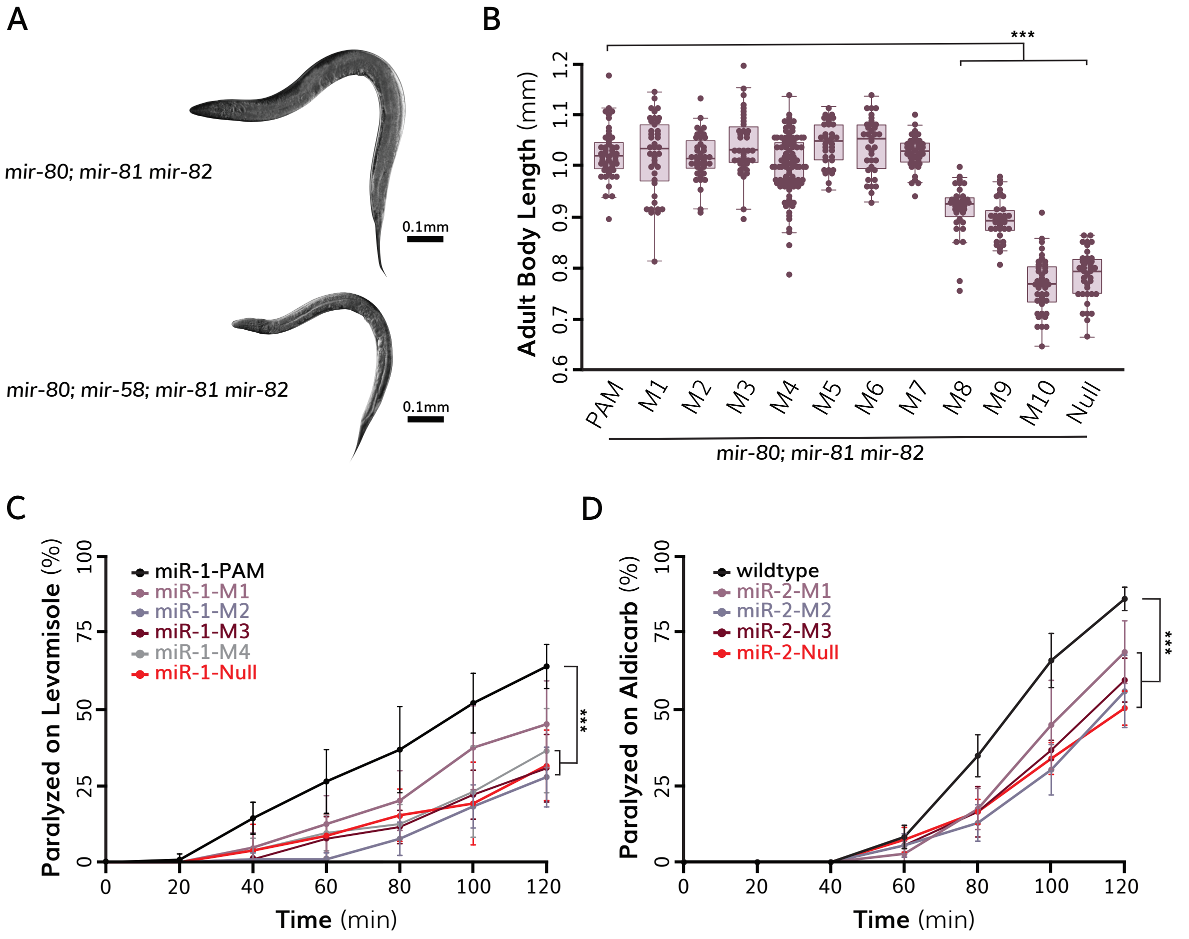


**Fig S5.** Phenotypic analysis of miRNA mutants. (A) Representative images of a (top) *mir-80; mir-81 mir-82* triple miR-58 family mutant and (bottom) *mir-80; mir-58; mir-81 mir-82* quadruple family mutant illustrates reduced adult body size associated with loss of miR-58 family function. (B) Quantification of adult body length in miR-58 mutants in the *mir-80; mir-81 mir-82* genetic background. Each dot represents one animal. Boxes range from the first to third quartile of the data. The thick bar represents the statistical median. Lines extend to the minimum and maximum data point excluding outliers that were defined as 1.5 times the interquartile range. (C) Percent of miR-1 mutant animals paralyzed on plates containing 200 µM aldicarb over time. Data are presented as mean ± standard deviation. (D) Percent of miR-2 mutant animals paralyzed on plates containing 1 mM aldicarb over time. Data are presented as mean ± standard deviation. (B-D) Statistical analysis: ***p<0.001.

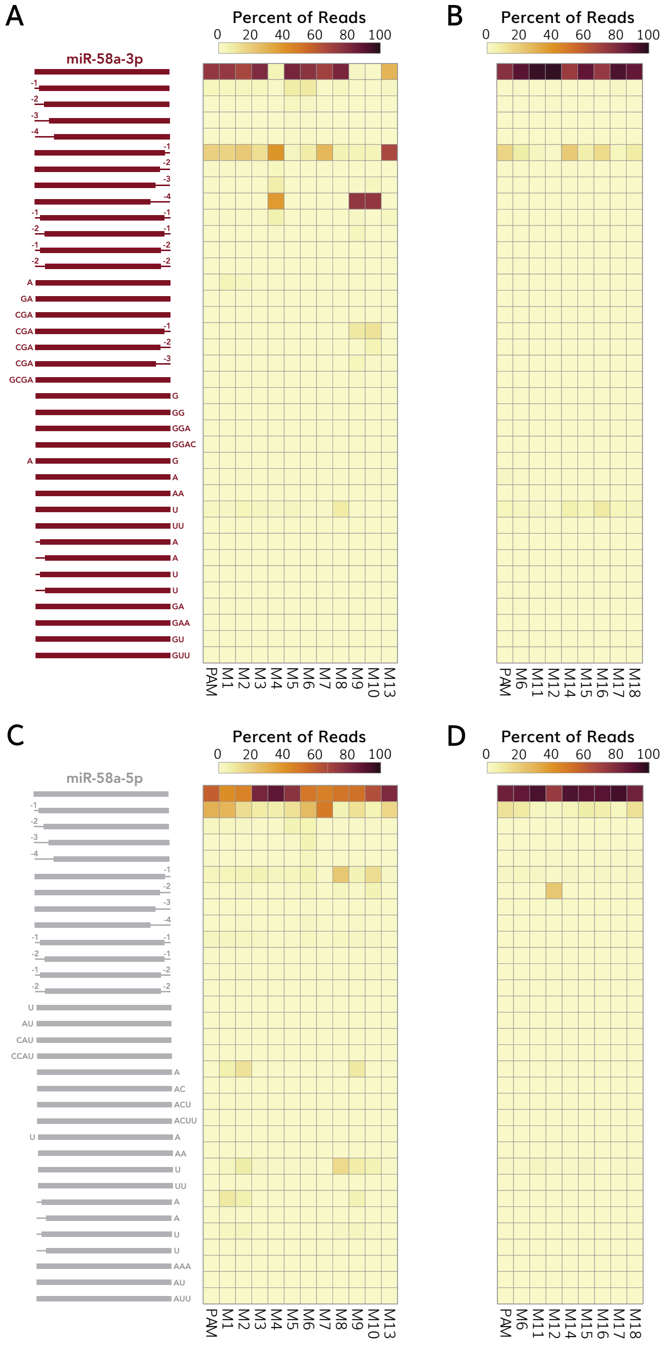


**Fig S6.** Analysis of miR-58 isomiRs in miR-58 mutants. (A-B) Heatmaps illustrating relative distribution of miR-58-3p (canonical guide strand) isomiRs in miR-58 mutants. Data are from small RNA sequencing data of L4-staged mutants that were created using the (A) Nextflex v3 or (B) Nextflex v4 small RNA sequencing kits. (C-D) Heatmaps illustrating relative distribution of miR-58-5p (canonical passenger strand) isomiRs in miR-58 mutants. Data are from small RNA sequencing data of L4-staged mutants that were created using the (C) Nextflex v3 or (D) Nextflex v4 small RNA sequencing kits. (A-D) Data are presented as the percent occurrence of a given isomiR within each individual mutant and do not reflect possible changes in miRNA abundance. Each isomiR is illustrated on the left. The maroon (guide) and gray (passenger) bars indicates the expected miR-58 sequence. Truncated isomiRs are indicated by negative values corresponding to the length of the truncation. Nucleotide extensions are depicted by the corresponding letters to each duplex end, which may include templated or non-templated (uridylation and adenylation) extensions.

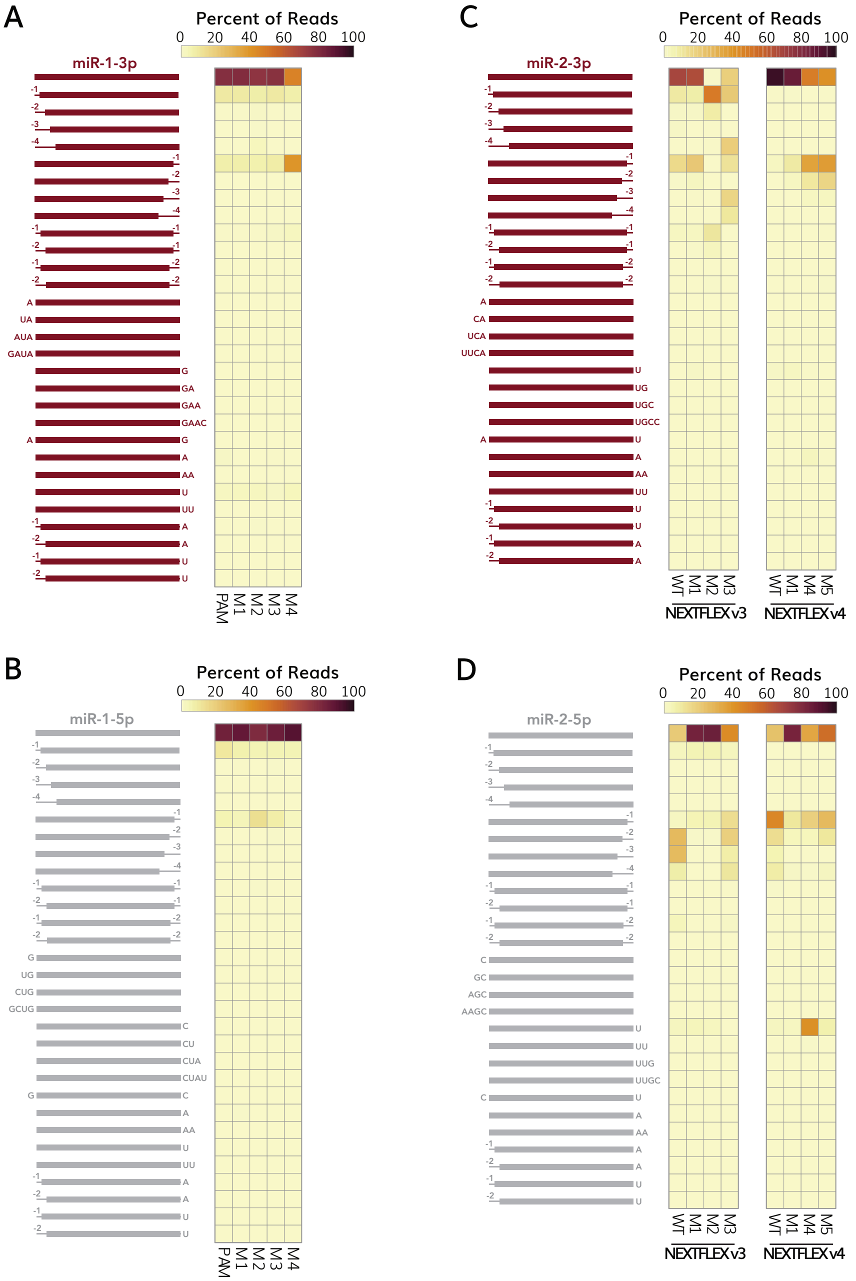


**Fig S7.** Analysis of miR-1 and miR-2 isomiRs in miRNA mutants. Heatmaps illustrate relative distribution of (A) miR-1-3p (canonical guide strand) or (B) miR-1-5p (canonical passenger strand isomiRs in miR-1 mutants. (A-B) Data are from small RNA sequencing data of L4-staged mutants. Heatmaps depict (C) miR-2-3p (canonical guide strand) and (D) miR-2-5p (canonical passenger strand) isomiR distribution in miR-2 mutants. (C-D) Data are from small RNA sequencing data of L4-staged mutants that were created using the (left) Nextflex v3 or (right) Nextflex v4 small RNA sequencing kits. (A-D) Data are presented as the percent occurrence of a given isomiR within each individual mutant and do not reflect possible changes in miRNA abundance. Each isomiR is illustrated on the left. The maroon (guide) and gray (passenger) bars indicates the expected miRNA sequence. Truncated isomiRs are indicated by negative values corresponding to the length of the truncation. Nucleotide extensions are depicted by the corresponding letters to each duplex end, which may include templated or non-templated (uridylation and adenylation) extensions. Note that pre-miR-2 contains a templated uracil immediately downstream of the 3' nucleotide of the mature miR-2 guide and passenger strands, preventing disambiguation of non-templated or templated addition of uracil to either strand.

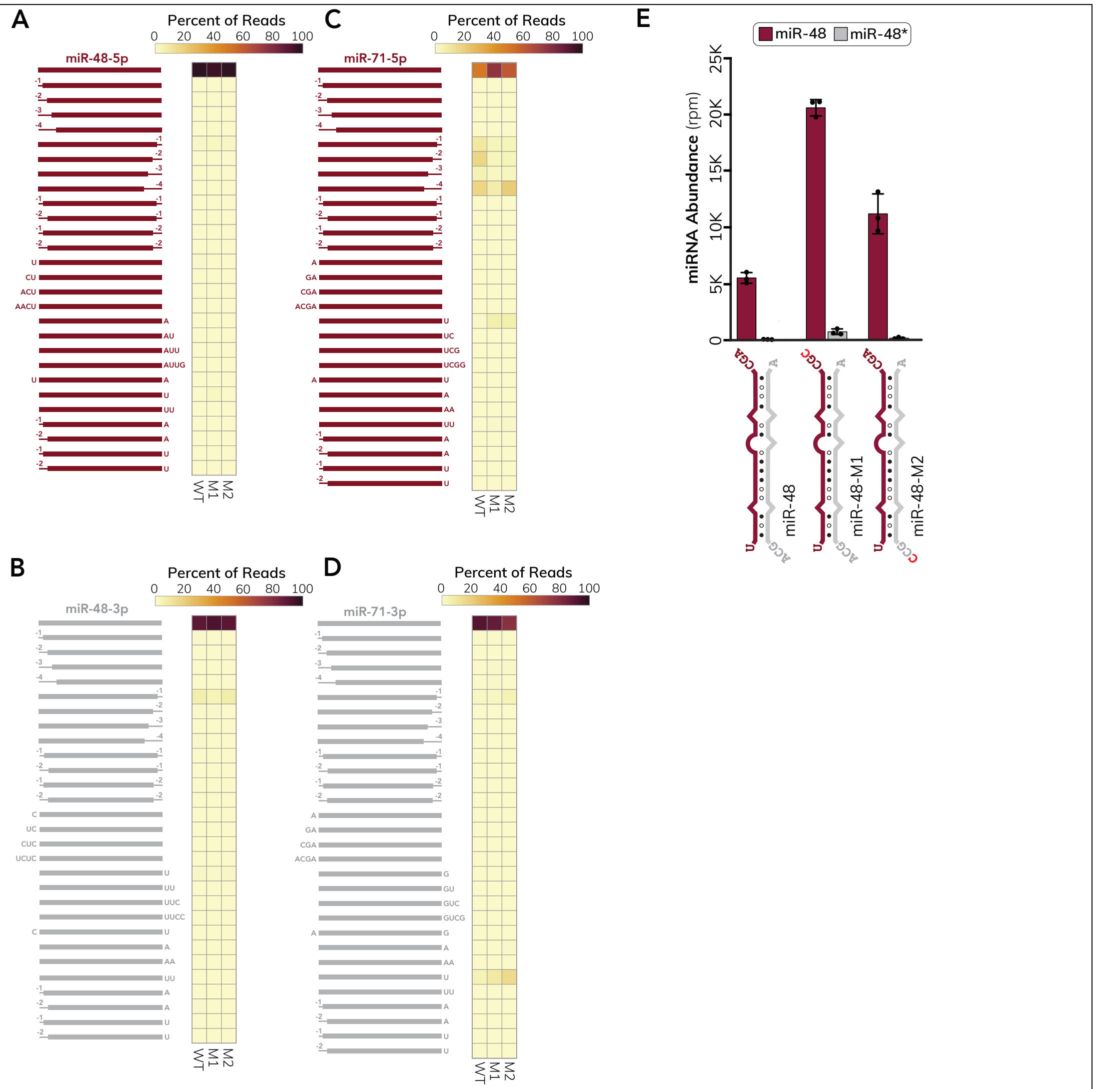


**Fig S8.** Analysis of miR-48 and miR-71 isomiRs in miRNA mutants. Heatmaps illustrate relative distribution of (A) miR-48-5p (canonical guide strand), (B) miR-48-3p (canonical passenger strand), (C) miR-71-5p (canonical guide strand) and (D) miR-71-3p (canonical passenger strand) in miR-48 or miR-71 mutants. (A-D) Data are presented as the percent occurrence of a given isomiR within each individual mutant and do not reflect possible changes in miRNA abundance. Each isomiR is illustrated on the left. The maroon (guide) and gray (passenger) bars indicates the expected miRNA sequence. Truncated isomiRs are indicated by negative values corresponding to the length of the truncation. Nucleotide extensions are depicted by the corresponding letters to each duplex end, which may include templated or non-templated (uridylation and adenylation) extensions. Note that pre-miR-48 contains a templated adenine immediately downstream of the 3' nucleotide of the mature miR-48 guide strand and a templated 3' uracil downstream immediately downstream of passenger strand, preventing disambiguation of non-templated or templated additions. Similarly, pre-miR-71 contains a templated uracil immediately downstream of the 3' nucleotide of the mature miR-71 guide strand. (E) (Bottom) Schematic representations of miR-48 duplex mutations that affect 3' nucleotide identity. The guide strand is shown in maroon, and the passenger strand is shown in gray. The nucleotide composition of dinucleotide overhangs are shown for each strand. Mutated nucleotides are indicated by red text. U:A pairs are illustrated by an open circle, G:U wobble pairs are indicated by an open circle with a vertical line and G:C pairs are represented by closed circles. RNAcofold was used to predict duplex structures at 20°C. (Top) Quantification of small RNA sequencing results for miR-48 mutants. Data are presented as mean ± standard deviation, with each dot representing quantification from a single library using the NEXTFLEX v4 Small RNA Sequencing Kit. Maroon bars correspond to miRNA guide strands and gray bars represent passenger strands. Normalization was performed by dividing raw read counts by total mapped miRNA reads and multiplying by one million to acquire reads per million (rpm) values.

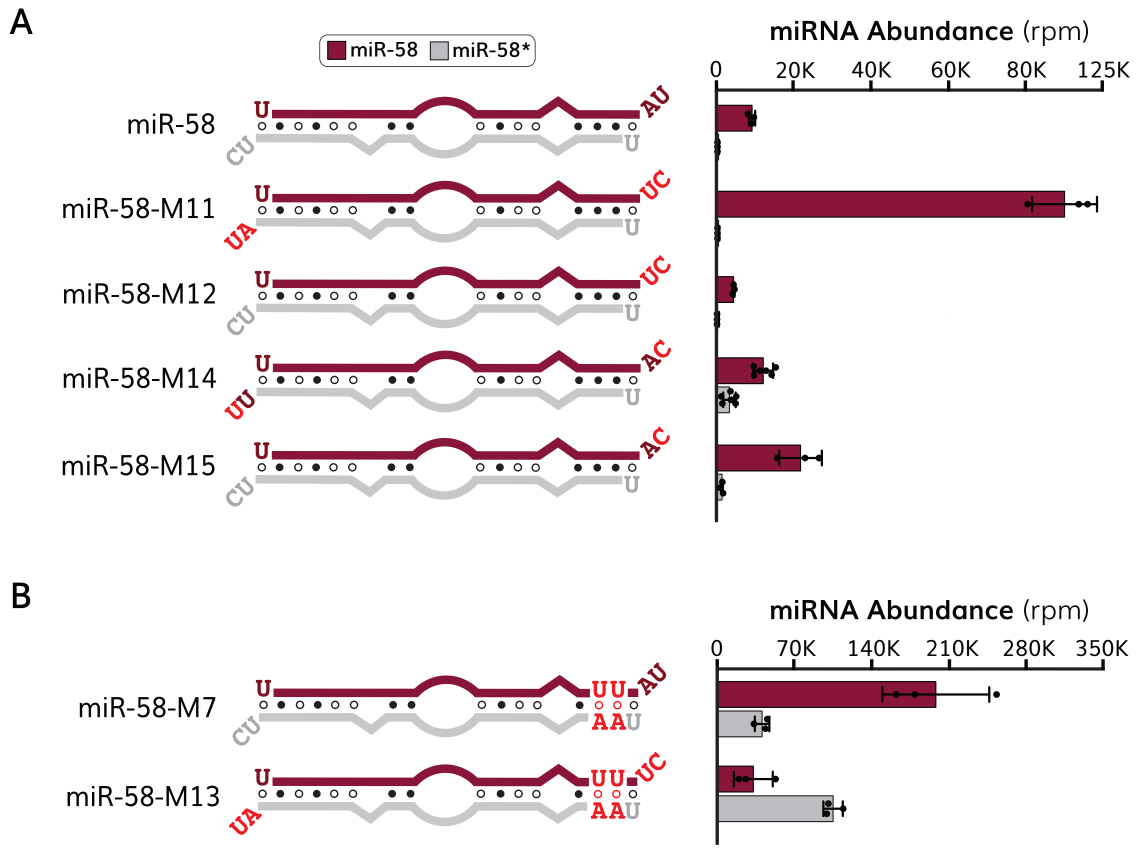


**Fig S9.** The composition of dinucleotide overhangs may influence miRNA strand selection. (A-B) (Left) Schematic representations of miR-58 duplex mutations that affect the nucleotide composition of dinucleotide overhangs. Mutations were made in a (A) wildtype miR-58 duplex background or (B) the miR-58-M7 mutant duplex. The guide strand is shown in maroon, and the passenger strand is shown in gray. Mutated nucleotides are indicated by red text. U:A pairs are illustrated by an open circle, G:U wobble pairs are indicated by an open circle with a vertical line and G:C pairs are represented by closed circles. RNAcofold was used to predict duplex structures at 20°C. (Right) Quantification of small RNA sequencing results for each miR-58 duplex strand. Data are presented as mean ± standard deviation, with each dot representing quantification from a single library using the NEXTFLEX v3 Small RNA Sequencing Kit. Maroon bars correspond to miRNA guide strands and gray bars represent miRNA passenger strands. Normalization was performed by dividing raw read counts by total mapped miRNA reads and multiplying by one million to acquire reads per million (rpm) values.

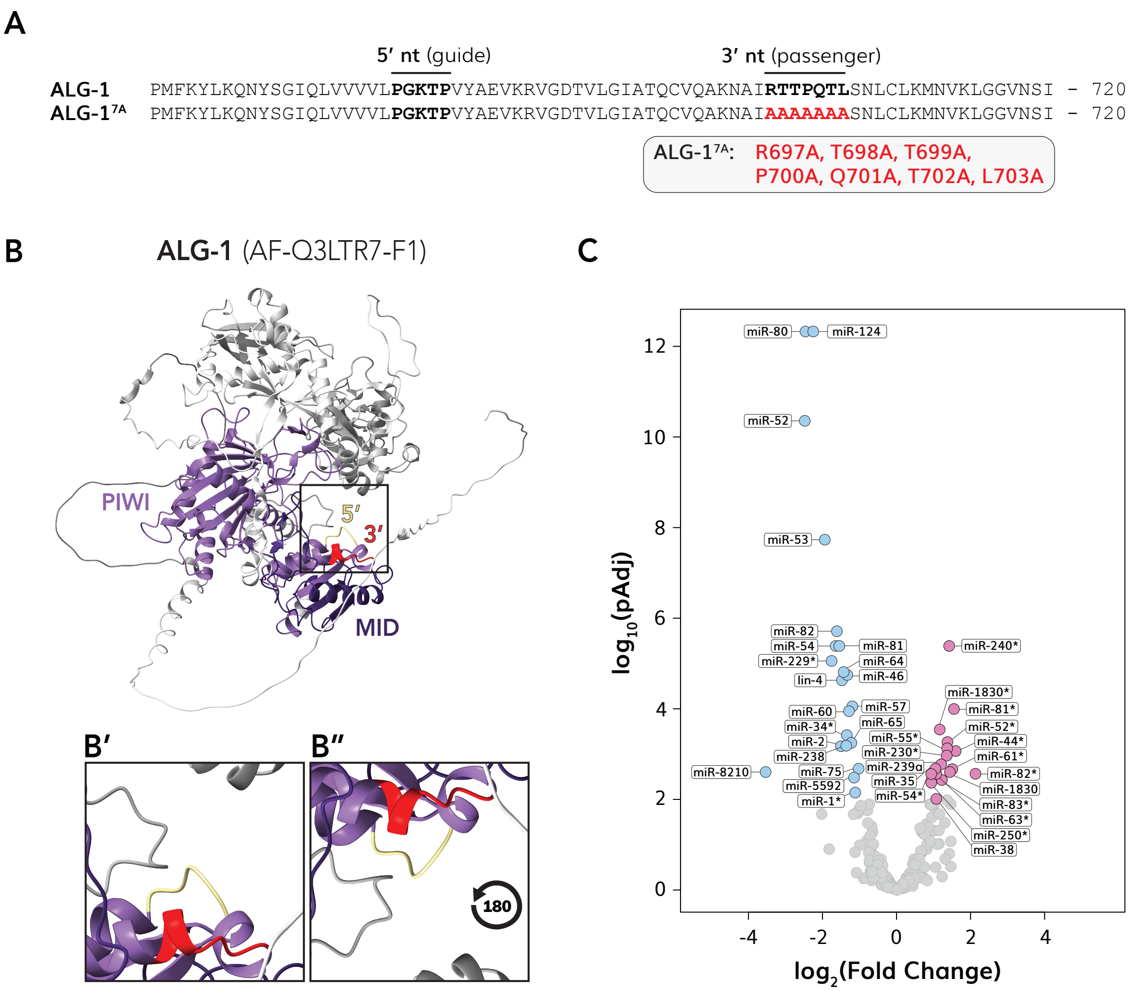


**Fig S10.** Functional analysis of ALG-1$^{7A}$ mutants. (A) Location of alanine substitutions in ALG-1$^{7A}$ mutants. Mutations are indicated by red text. The location of the ALG-1 5' nucleotide binding pocket(*13*, *14*) and putative 3' nucleotide binding pocket(*17*) are bolded. (B) Physical location of 5' nucleotide binding pocket (yellow) and putative 3' nucleotide binding pocket (red) on the Alphafold predicted structure of ALG-1 (AF-Q3LTR7-F1). The MID (dark purple) and PIWI (light purple) domains are marked. ChimeraX v1.9 was used for structure visualization. Inset highlights relative position of the 5' nucleotide binding pocket and putative 3' nucleotide binding pocket within the RNA-binding groove of ALG-1 (B') and rotated 180° (B"). (C) Differential expression analysis of ALG-1$^{7A}$ mutants compared to wildtype (N2) controls was performed using the 'DESeq2' package for R. Each dot represents one miRNA strand. Statistical significance was considered at p<0.01. Upregulated miRNAs are labelled and colored pink while downregulated miRNAs are labelled and colored blue.

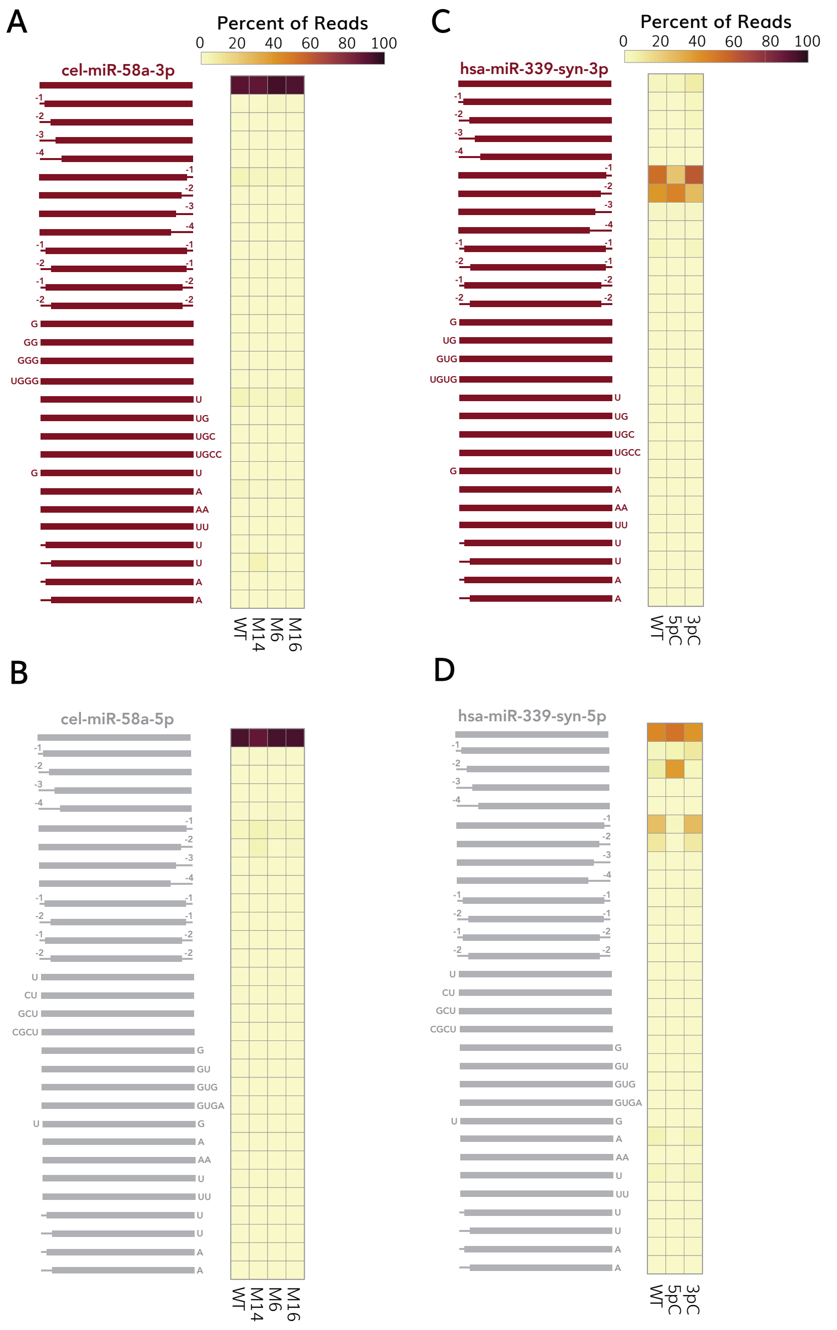


**Fig S11.** Analysis of isomiRs from miRNAs transfected into human HEK293T cells. Heatmaps illustrate relative distribution of (A) miR-58a-3p (*C. elegans* guide strand), (B) miR-58a-5p (*C. elegans* passenger strand), (C) miR-339-syn-3p (expected guide strand) and (D) miR-339-syn-5p (expected passenger strand) in miR-58 or miR-339-syn mutants. The miR-339-syn mutants carry an additional internal nucleotide substitution that is not expected to affected duplex structure (Supplementary Table 15), which allowed us to distinguish exogenous and endogenous miR-339 variants. (A-D) Data are presented as the percent occurrence of a given isomiR within each individual mutant and do not reflect possible changes in miRNA abundance. Each isomiR is illustrated on the left. The maroon (guide) and gray (passenger) bars indicates the expected miRNA sequence. Truncated isomiRs are indicated by negative values corresponding to the length of the truncation. Nucleotide extensions are depicted by the corresponding letters to each duplex end, which may include templated or non-templated (uridylation and adenylation) extensions. Note that the Ultramir scaffold contains a templated uracil immediately downstream of the 3' nucleotide of the 3p, preventing disambiguation of non-templated or templated uridylation for the 3p strand.

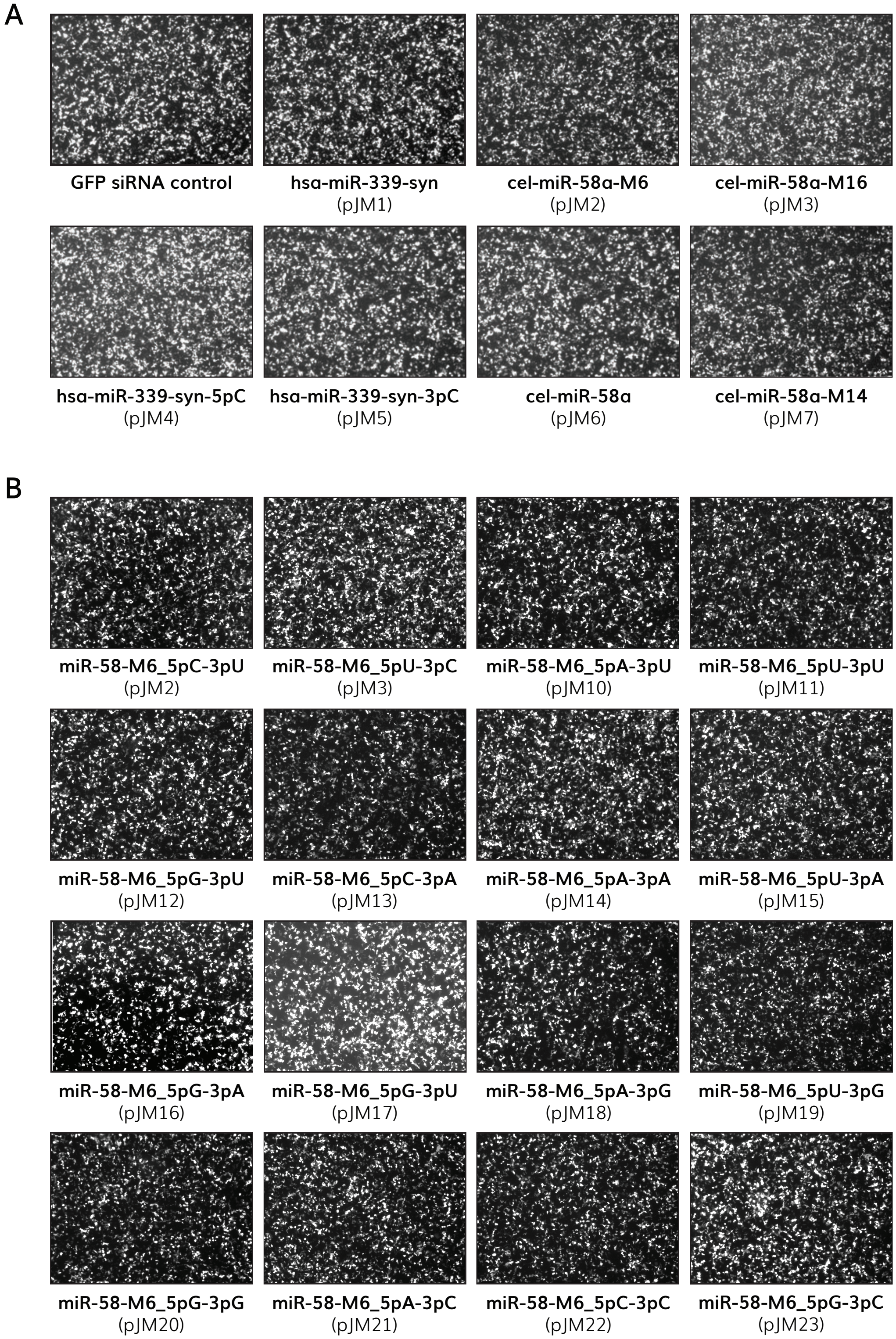


**Fig S12.** ZsGreen expression in transfected human HEK293T cells. Representative images of HEK293T cells expressing ZsGreen prior to harvesting. (A) Images of samples corresponding to Figure 6B. (B) Images of samples corresponding to Fig 6C.
